# Phosphodiesterase type 1 regulates adenosine 2A receptor-associated cAMP signaling at the plasma membrane to increase myocardial contractility

**DOI:** 10.1101/2025.10.03.680406

**Authors:** Michael L. Fitch, Maria Alexandria Martins Goncalves, Mary Tang, Evan D. Kelly, Emily C. Cheung, Purvaj Kandula, Emily Jones, Szymon Sornat, Orien Sun, Manuela Zaccolo, Matthew W. Kay, Grace K. Muller

## Abstract

**BACKGROUND:** Phosphodiesterase type 1 (PDE1) inhibition exerts inodilatory effects in pre-clinical models and human heart failure patients with reduced ejection fraction (HFrEF). PDE1 hydrolyzes cyclic nucleotide cAMP in a soluble but not microsomal fraction of the human myocardium. PDE1 may exert domain-specific effects, but the mechanism whereby PDE1 compartmentalization induces an inotropic change remains unknown. We sought to elucidate PDE1 regulation of cAMP and contractility.

**METHODS:** Pharmacologic modulators of PDEs or G-protein coupled receptors (GPCRs) were used to study PDE1 signaling mechanisms in healthy and failing guinea pig hearts. Tissue fractionation examined the localization of different PDE types. Live cell imaging experiments assessed cytosolic and sarcolemmal membrane cAMP level ([cAMP]) and protein kinase A (PKA) activity changes in cells transduced with Förster resonance energy transfer (FRET) biosensors. Sarcomere length and intracellular calcium changes monitored contractile changes in electrically paced cells. Coronary flow and left ventricular (LV) developed pressure were measured in ex vivo Langendorff perfusion heart studies.

**RESULTS:** PDE1 isoforms were found not only in the soluble, but also in the microsomal fractions of the guinea pig heart. PDE1 hydrolysis of cAMP was greater at the sarcolemma compared to the cytosol. PDE1 specifically regulated a pool of cAMP associated with the G⍺_s_ protein coupled receptor adenosine 2A receptor (A_2A_R) at the sarcolemma, without activating PKA. A_2A_R/PDE1 regulation induced positive inotropic and lusitropic changes in healthy and failing guinea pig cardiomyocytes and in the myocardium ex vivo.

**CONCLUSIONS:** PDE1 is the major regulator of cAMP pools generated by A_2A_R activation at the sarcolemma. Functionally, this regulation induces inotropic and lusitropic effects in cardiomyocytes and at the whole organ level. Thus, PDE1 is compartmentalized at the membrane with A_2A_R, and this regulation determines cardiomyocyte contractility in healthy and failing hearts.

## Introduction

Affecting 0.8% of the world population, heart failure (HF) is a global burden especially among the elderly.^1^ Acute decompensated HF is the leading diagnosis for hospital admissions, in which patients present with serious new or worsening HF symptoms and signs.^2^ Current treatments include diuretics, vasodilators, and inotropes such as dobutamine and milrinone. Dobutamine works as a β-adrenergic receptor agonist to stimulate heart contractility, increasing levels of the cyclic nucleotide cAMP. Milrinone similarly increases cAMP by inhibiting phosphodiesterase type 3 (PDE3).

PDE3 is one of the two key enzymes that hydrolyze cAMP in large mammalian hearts, including humans.^3^ The buildup of cAMP activates protein kinase A (PKA), which will phosphorylate Ca^2+^-handling proteins and myofilament proteins to enhance myocyte contractility and relaxation. Inhibition of the other key cAMP-hydrolyzing enzyme, PDE type 1 (PDE1),^3,4^ also induces vasodilatory effects in animal models^5^ and human patients of HF with reduced ejection fraction (HFrEF).^6^ In a Phase Ib/IIa trial, the PDE1 inhibitor lenrispodun worked as an inotrope to improve LV contractility and decrease systemic vascular resistance in HFrEF patients.^6^

Increasing body of evidence indicates that cell signaling pathways are spatially regulated. Also referred to as nanodomains,^7^ compartmentalization describes how a common second messenger such as cAMP can exert a specific cellular response based on its spatiotemporal distinction.^8,9^ At the sarcoplasmic reticulum (SR), PDE3A modulates the PKA phosphorylation level of phospholamban (PLN) and thus Ca^2+^ reuptake.^10,11^ PDE4D regulates ryanodine receptor function and Ca^2+^ release,^12^ though the enzyme plays a much lesser role in the human heart. PDE3A is also expressed in the nucleus, where its regulation of the TGF-β/Smad4 signaling pathway controls hypertrophic response via modulation of histone deacetylase 1 enzymatic activity.^13^ In the human myocardium, PDE3 accounts for 69% of the microsomal cAMP hydrolysis. PDE1, on the other hand, accounts for 78% of the soluble cAMP hydrolysis.^4^

Uniquely activated by Ca^2+^/calmodulin, PDE1 has isoform-dependent substrate specificity. PDE1A hydrolyzes cGMP, while PDE1C hydrolyzes both cGMP and cAMP with similar affinity.^4^ PDE1A expression is prominent in rats and mice; PDE1C expression is greater in guinea pigs and larger mammalian hearts, including humans’.^5^ PDE1C forms a multi-protein complex with transient receptor potential canonical channel 3 (TRPC3) and the GPCR adenosine 2A receptor (A_2A_R). Functionally, ARs are important cardioprotectors during ischemic injury,^14^ where the ligand adenosine levels increase in damaged areas. Extracellular cAMP efflux, downstream of A_2A_R activation, is cardioprotective in oxidative stress by exerting an anti-apoptotic effect.^15^ Similarly, PDE1 inhibition (PDE1i) or genetic ablation is anti-apoptotic,-fibrotic, and - hypertrophic.^16^ More acutely, A_2A_R activation induces inotropic effects. However, this is debated,^17^ and has not been considered in the context of concomitant PDE1i. The nanodomain in which PDE1 modulates cAMP/PKA signaling has not been elucidated.

The compartment differences between PDE1 and PDE3 activity may be key to understanding a clinically relevant question in consideration of the use of either inhibitor. The use of PDE3 inhibitors is not recommended as long-term HF treatment because of fatal arrhythmias and reduced survival in dog models and human patients with HFrEF.^9^ At the cellular level, PDE1i is a less potent inotrope compared to PDE3i, possibly because PDE3i actively increases SR Ca^2+^ reuptake, while PDE1i does not. And yet, both PDEs, when inhibited, increase the voltage-activated plasma membrane Ca^2+^ channel Ca_v_1.2 (L-type Ca^2+^ channel) in a PKA-dependent manner.^18^ Therefore, we hypothesized that PDE1 regulation of the cAMP signaling pathway at the sarcolemmal membrane may be an important determinant of cardiac contractile function. We tested this hypothesis using in vitro, ex vivo, and in vivo techniques.

## Materials and Methods

### Data Availability

The data that support the findings of this study are available from the corresponding author upon reasonable request. All reagents used in this study are available commercially, except for IC200014 that was provided from Intracellular Therapies, Inc. under a research agreement. All other reagents are commercially available, and listed in the Major Resource Table in the Data Supplement.

### Animals

Hartley guinea pigs were purchased from Charles River Laboratories or Hilltop Lab Animals. Animal care and procedures were performed at Loyola University of Chicago or George Washington University, in facilities accredited by the Association for the Assessment and Accreditation of Laboratory Animal Care International. Protocols pertaining to the present study were approved by the Institutional Animal Care and Use Committees at respective institutions where the procedures were performed, and were in accordance with the Guide for the Care and Use of Laboratory Animals published by the US National Institutes of Health.

### Guinea Pig Aortic Constriction Surgery

A modified aortic constriction (AC) surgery was performed on guinea pigs from an original model established by Liu et al.^19^ Male guinea pigs weighing 280 to 400 g were anesthetized with isoflurane (4% to induce and 1.5-2% to maintain) and mechanically ventilated. Aseptic techniques were applied to perform thoracotomy, cutting along the sagittal plane between the 4^th^ and 5^th^ intercostal space. A suture was tied around the ascending aorta with a 14G needle used as a spacer. The needle was subsequently removed. The pectoral muscles and the skin were closed using absorbable and non-absorbable sutures, respectively. In sham animals, muscles and skin layer were closed without the constriction procedure. Guinea pigs were allowed to recuperate post-surgery. Post-surgery management included daily monitoring for a week with metoclopramide (0.5 mg/kg q 12 hours) and buprenorphine slow release (0.3 mg/kg q 24 hours) injections for the first three days. In a cohort of 19, the mortality rate due to surgical complication was 10.5% (2/19). One animal died following recovery to an unknown cause, and 4 were removed from study because the surgery failed to induce HF during study period.

### Echocardiography and Analysis

Baseline echocardiography was performed in anesthetized guinea pigs (4% isoflurane to induce and 1.5-2% to maintain) in the week prior to surgery, then serially every 2 weeks post AC surgery, up to 14 weeks post-surgery. An MX250 probe (FUJIFILM) was used to image the left ventricle in the parasternal long-axis view in Brightness Mode, followed by Motion-Mode in the short-axis view. Using M-mode analysis in Vevo Lab software (FUJIFILM), the following parameters were obtained: heart rate, LV ejection fraction (EF), fractional shortening (FS), anterior/posterior wall thickness (AWd or PWd) and interior diameter (LVd) in diastole. The LV weight was calculated using the following equation: 0.8 x 1.04 x ((AWd + LVd + PWd)^3^ – LVd^3^) + 0.6g. Animals were euthanized for heart tissue removal when the left ventricular EF reached ≤40% guinea pigs in the AC group. Body weight-matched sham animals served as controls.

### Thoracotomy Retrieval of the Hearts

Guinea pigs, ranging from 300 g to 500 g (≃2-3 months old male and female for healthy animals) or 495 g to 810 g (≃5-7 months old male for sham or AC animals), were sedated with medetomidine (0.75 mg/kg) and ketamine (90 mg/kg). For those used in excised perfused heart studies, guinea pigs were anesthetized with a ketamine/xylazine (50+5 mg/kg) cocktail followed by isoflurane (3-5%) inhalation until cessation of pedal reflex. Hearts were then rapidly excised after thoracotomy and processed for each experiment, either separated into chambers and snap frozen in liquid nitrogen for biochemical analysis, or immediately prepared for an enzymatic isolation of cardiomyocytes, or prepared for ex vivo isovolumic working heart studies, as described below.

### Fractionation of Ventricular Heart Tissue

A serial centrifugation protocol was applied to subdivide frozen heart ventricular tissue into fractions enriched in nuclei, cytosol, and microsomes, as adapted from published protocols.^4,20^ A rotor-stator system (gentleMACS Octo Dissociator, Miltenyi Biotec) was used to homogenize frozen ventricular tissue in 3x 15s cycles. The homogenization buffer (2ml / tissue weight g) consisted of [in mM: 10 PIPES, 2 EDTA, 2 EGTA, 250 sucrose, pH 7.0], and was supplemented with 1x protease inhibitor cocktail (cOmplete Mini, Roche). Sequential centrifugation steps were performed at 4℃, beginning with the isolation of the nuclei: 1,000 x g for 5 min. The pellet was dissolved in 2x volume of homogenization buffer and the centrifugation step was repeated to yield the nuclei fraction. The pooled supernatant was ultracentrifuged at 30,000 x g for 1 hour to yield a supernatant consisting of soluble proteins (cytosol). The pellet was dissolved in 2x volume of homogenization buffer supplemented with 0.6 M KCl, and ultracentrifuged at 100,000 x g for 1 hour. The supernatant was discarded, and the pellet represented the microsomal fraction, consisting of plasma membrane and organelle. Pellets of interest were reconstituted in a storage buffer [in mM: 300 sucrose, 5 HEPES, pH 7.4] and assayed for protein content using Bradford Assay (Bio-Rad), and stored at-80℃ until ready for western blotting.

### Isovolumic Working Heart Studies

Excised hearts were placed in heparinized, ice-cold modified Krebs-Hennseleit (KH) perfusate consisting of [in mM: 115.0 NaCl, 3.3 KCl, 2.0 CaCl_2_, 1.4 MgSO_4_, 1.0 KH_2_PO_4_, 25 NaHCO_3_, 6 glucose, pH 7.4]. To retrogradely perfuse the heart, the aorta was cannulated with a short 14G catheter, secured with a 6-0 suture, and flushed with 10 mL ice-cold perfusate to remove residual blood. Hearts were then Langendorff perfused at a constant hydrostatic pressure of 75 mmHg with perfusate bubbled with 95% O_2_, 5% CO_2_ and maintained at 37 ± 0.5℃. Hearts were partially submerged in perfusate and positioned with the LV posterior wall facing up. The left atrial appendage was removed and the mitral valve was vented. Aortic perfusate temperature, the electrocardiogram, and coronary flow were continuously acquired and monitored using Labchart (ADInstruments). After a 20 minute stabilization period, a deflated latex balloon (size 5, Harvard Apparatus) connected to a pressure transducer (World Precision Instruments, Sarasota, Fl, USA) was inserted in the LV and the heart was acclimated for 10 minutes. The balloon was then inflated with water to a diastolic pressure of 8 mmHg, and heart rate and LV developed pressure (LVDP) were monitored continuously using Labchart.

Contractility (dP/dt _max_) and relaxation (dP/dt _min_) were calculated using the derivative of LVDP. Hearts were paced at 4.58 Hz (275 BPM) from an electrode placed on the posterior right ventricle. Hearts that did not maintain consistent pacing at 275 BPM were excluded from analysis. Each heart was randomly assigned to three treatment groups: 1) 300nM (carboxyethylphenethyl-aminoethyl-carboxamido-adenosine 21680 (CGS) alone, 2) 1nM IC200041 (2000) alone or 3) priming with 1nM 2000 for ten minutes followed by 300nM CGS. These drugs were administered into aortic flow at a rate that matched the coronary flow. The response to drug administration was recorded for ten minutes.

### Adult Guinea Pig Cardiomyocyte Isolation

To isolate cardiomyocytes, excised hearts were submerged in ice-cold Tyrode’s [in mM: 140 NaCl, 5 KCl, 1 MgCl_2_, 10 HEPES, 5.5 glucose, pH 7.5 at 37℃] supplemented with 2.8 au/ml heparin. The aorta was cannulated and hung on a customized Langendorff perfusion system (Radnoti), consisting of a water-jacketed coiled heat exchanger, bubble trap, and collecting chamber that encased the heart. Perfusate was maintained at 37℃ using a water circulating pump (Thermo Scientific Haake SAHARA S7). Blood was rinsed out by perfusing the heart with Tyrode’s for 5 mins using a peristaltic pump (Masterflex) set at 8.7 ml/min, then digested for 6 mins with a digestion solution (18 mg collagenase type II (Worthington) and 3 mg protease type XIV (Sigma) in 40 ml of the Tyrode’s. The perfusate was switched to a high potassium solution [in mM: 120 KH_2_PO_4_, 25 KCl, 1 MgCl_2_, 10 HEPES, 1 EGTA, 5.5 glucose, pH 7.4 at 37℃] for 5 mins. Upon completion, the ventricles were removed from the heart, and mechanically triturated first with scissors and next with a blunt-ended transfer pipette. The dissociated tissue was filtered through a 200-micron mesh and allowed to settle. The solution was replaced 2-4 times with M199 [in mM, 5 taurine, 2 l-carnitine, 5 creatine, 0.0001 insulin, 2 mg/ml BSA, 1x pen/strep] to enrich for living cardiomyocytes.^21^ Cardiomyocytes were used immediately for biochemical or functional experiments, or cultured to perform live cell imaging experiments, as described below.

### FRET Imaging and Analysis

Freshly isolated guinea pig cardiomyocytes were plated (5-9 x 10^4^ cells) on 25 mm coverslips pre-coated with laminin (diluted 1:50 in phosphate buffered saline (PBS)) for at least two hours. Cells were kept in a 37℃ incubator with 5% CO_2_. Cardiomyocytes were transduced with adenovirus encoding FRET biosensors Epac-S^H187^ (H187),^22^ plasma membrane-Epac-S^H187^ (pm-H187)^23^ or AKAR4^24^ at 1000 MOI. After 3-4 hour of incubation, the media containing adenovirus was replaced with fresh M199 media as previously reported.^8^ FRET experiments were performed within 1-2 days, with optimal imaging times for AKAR4 occurring between 20-30 hours, and H187 and pm-H187 occurring 24-48 hours post-infection.

FRET experiments were performed using an Eclipse Ti2 microscope (Nikon) customized with an ORCA-Fusion CMOS camera (Hamamatsu), a motorized stage, a filter wheel, and a LED light source (Spectra III, Lumencor). Cardiomyocytes were sequentially imaged every five seconds using a common 440nm excitation, with emission collected for cyan (476/27 nm) and yellow (544/24 nm). At the time of experimentation, coverslips were transferred to an Attofluor chamber in 1.8mM Ca^2+^ Tyrode’s and visualized using a 40x oil objective. Experiments started typically within 3 minutes, and after an additional 3 minutes of baseline recording, cells were treated with pharmacologic agents as described in text, including non-discriminatory basal adenylyl cyclase activator forskolin (fsk), adenosine (Ado), adenosine A_2A_ receptor agonist

CGS21680 (CGS), or adenosine A_2B_ receptor agonist Bay 60-6583 (BAY) prior to a PDE inhibitor (type 1 inhibitor: IC200041; type 3 inhibitor: cilostamide, type 4 inhibitor: rolipram). To assess the role of A_1_R, some coverslips were pretreated with its specific antagonist dipropylcyclopentylxanthine (DPCPX) for 35-45 minutes. Every experiment was concluded with a pan-phosphodiesterase inhibitor 3-Isobutyl-1-methyxanthine (IBMX) and fsk to induce maximal [cAMP].

Data analysis was performed offline. Regions of interests were background-subtracted and exported as a ratio of CFP/YFP (H187 and pm-H187) or YFP/CFP (AKAR4) using Elements software (Nikon). The data were normalized for baseline and maximal response. The baseline (0%) was considered as the mean of the ratio during the 15-seconds preceding the first drug introduction, whereas the mean of the peak response following IBMX+fsk treatment was considered as 100%. Where a second drug was introduced during the experiment, the steady state FRET change (%) of the drug in relation to the prior steady state level was considered as the drug response. These data were plotted against time using Prism 10.0 (GraphPad).

#### Cardiomyocyte Sarcomere Shortening and Intracellular Calcium Measurements

Cardiomyocyte contractility and intracellular Ca^2+^ measurements were performed using an inverse microscope (TE-2000U, Nikon) customized with MyoCam2 (IonOptix) and photon multiplier tubes (PMTs). Brightfield light was provided by a halogen bulb light source (Nikon). All functional measurements were performed as reported before.^5,18^ Freshly isolated cardiomyocytes were loaded with 1 µM fura-2 AM (Invitrogen) for 12 min, followed by two washes of 15 minutes, each, with Tyrode’s. Cells were paced using an electrical pacer (MyoPacer) at 1 Hz for 2 milliseconds using 40 volts, and excited at 340/380 at a rate of 250 Hz (Cairn). Baseline was established for three minutes under these settings with continuous perfusion of 1.8 mM CaCl_2_ Tyrode’s solution at 2 ml/min. An initial time course determined the peak drug response time to be within 2-4 minutes. Subsequent experiments were conducted using MultiCell Lite, in which multiple cells are selected and monitored sequentially and repeatedly using a motorized stage (Prior). Sarcomere shortening (SS) and CaT recordings were analyzed for baseline and drug response using CytoSolver. Cells falling within mean±2SD for peak percent SS change were included for further analysis as described before.^18^ Parameters were analyzed for peak SS % change, time to 50% contraction, and tau; and for CaT peak change, time to 50% upstroke, and tau.

### Immunofluorescence and Imaging

Guinea pig isolated cardiomyocytes were plated at 10 x 10^4^ cells per coverslip pre-coated with laminin as described above. Cells were cultured in M199 for 1-2 days. Cells were fixed using 10% formalin for 10 min at RT. Following 3x rinses with PBS, cells were permeabilized with 0.1% Triton X-100 in PBS for 30 min, and blocked with 10% BSA (Sigma) in 0.1% Triton X-100 PBS for 1 hour, all at RT. Primary antibodies were incubated overnight at 4℃ at their indicated concentrations (Major Resources Table Data Supplements) in staining buffer [0.1% Triton X-100, 1% BSA in PBS]. After 3-5 washes in PBS, cells were incubated with appropriate secondary antibody for 1 hour at RT in staining buffer. Slides were mounted with ProLong (Invitrogen) and imaged using confocal microscope (SoRa Nikon). Where we sought to determine changes in phosphorylated PKA substrates, cells were treated with different drugs dissolved in Tyrode’s for 5 min at RT and washed twice in Tyrode’s supplemented with phosSTOP (Roche) before staining as already described. A customized code (deposited https://github.com/JamesSzczerkowski/Grace-Muller-Lab-2025.git) was developed to quantitate staining intensity across conditions.

### Western Blotting

Western blotting was performed as described.^18^ Freshly isolated cardiomyocytes were incubated with DMSO as negative control, or isoproterenol (ISO) or IBMX+fsk for positive controls, or various drugs in 1.8 mM Ca^2+^ Tyrode’s for 5 min rotating at RT. Cells were homogenized in lysis buffer (Cell Signaling) supplemented with a protease inhibitor cocktail (cOmplete Mini, Roche) and phosphatase inhibitor cocktail (PhosSTOP, Roche) on ice using a pipette, followed by passaging 7-10x via a 29G 1/2cc insulin syringe. Protein was quantified using Bradford Assay (Bio-Rad) and gel-loading samples were prepared using 4x Laemmli buffer (Bio-Rad) spiked with β-mercaptoethanol (Bio-Rad). Samples were allowed to reduce at RT for 20 min. Proteins (10-20 µg) were separated on gel electrophoresis at 90 V 10 minutes then 100V 1.5-2 h, followed by semi-dry transfer (Bio-Rad) onto 0.22 µm nitrocellulose membranes. To probe for phospholamban (PLN), proteins were run on a 12% gel; for all other proteins 4-15% gels were used. Membranes were blocked with Intercept Blocking Buffer TBS (Li-Cor). Antibody incubation was done in the same buffer supplemented with 0.2%

Tween for all proteins, with the exception of PDE1C, where membranes were blocked in and probed with antibody diluted in 0.1% KLP (SeraCare). Antibodies, their source and concentrations are in the Major Resource Table Data Supplements. Li-Cor Imager (Odyssey) was used to scan membrane, and Fiji^25^ was used to quantitate densitometry values.

## Statistical Analysis

Data are presented as mean ± SEM bar graphs or as box-whisker plots indicating the median and 5^th^-95^th^ percentile range. Male data are presented as black or empty, while female data are presented as red filled-in circles. For data with normal distribution, parametric analysis was performed. These included Student’s T-test, 1-way ANOVA, or 2-way ANOVA with indicated post-hoc tests. Nonparametric tests included Mann-Whitney U or Kruskal-Wallis tests. We considered p<0.05 as a statistically significant cutoff value, and all exact p-values are provided where significant.

## Results

### PDE1C is found in both the soluble and microsomal factions of the heart

To test the idea that PDE1C is expressed at the sarcolemma, we performed immunocytochemistry-based comparison between PDE1C and PDE3A in guinea pig ventricular cardiomyocytes. PDE1C staining pattern was diffused throughout the entire cell, along with at the intercalated disks (Suppl Fig 1A). Analysis of the micrographs showed that intracellularly, both PDEs localized with the interfilament protein desmin that marks sarcomere Z-disks (Suppl Fig 1). We also stained for PDE1C or PDE3A with the caveolae marker caveolin-3 within the same cell. As shown in Fig 1A, unlike PDE3A that showed an outlining of the sarcolemma (asterisks), detection of PDE1C at the membrane was restricted to the intercalated disc (arrows). A biochemical analysis of myocardial fractions showed that PDE1C is found not only in the cytosol but also in the Cav3-enriched microsomal (i.e., particulate) as well as the nuclei fraction (Fig 1B). This pattern was like that of PDE3A. The cGMP-hydrolyzing PDE1A was more notably enriched in the soluble, then the nuclei, and much less so in the microsome. PDE4D has several splice variants. We found some cytosolic bands likely corresponding to PDE4D1, D5, D7;^26^ and microsomal variants likely corresponding to PDE4D2, 8 and 9, as reported previously.^27^

**Fig 1.**
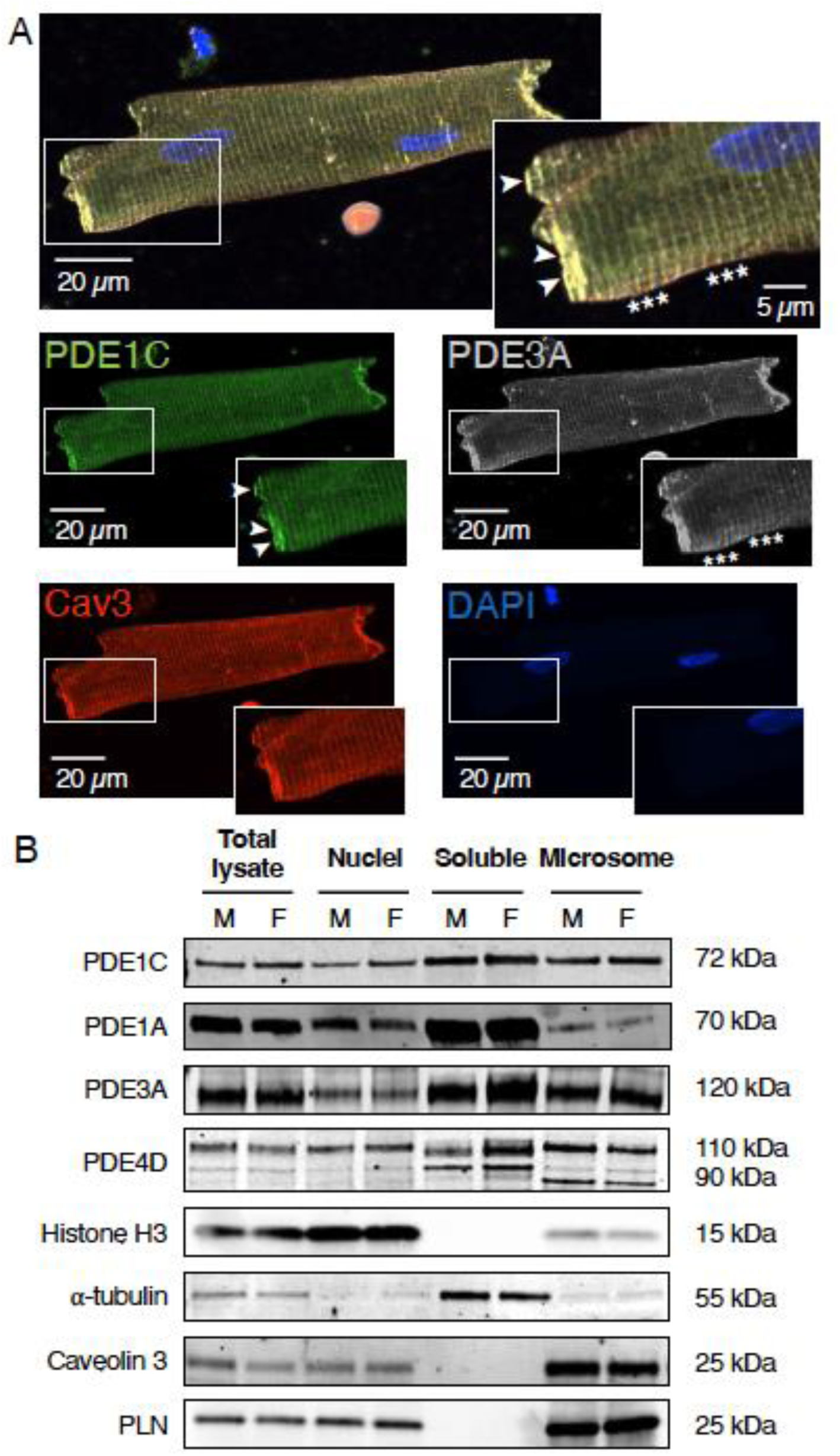
PDE1C is expressed in the soluble, microsomal and nuclei fraction of cardiomyocytes. A) Representative immunostaining of PDE1C and PDE3A in adult guinea pig ventricular cardiomyocytes co-stained with caveolin3 (Cav3) and DAPI. B) Representative western blot analysis of PDE1C, PDE1A, PDE3A and PDE4D in fractions of adult guinea pig ventricular heart tissue: histone H3 was used as a nuclear marker, ⍺-tubulin as a cytoplasm (soluble) fraction representative, and Cav3 and phospholamban (PLN) as microsomal. (N=6)

### Sarcolemmal PDE1 enzymatic activity is greater than in the cytosol

We tested if PDE1 may have a functional role to regulate [cAMP] at the membrane. To detect PDE activity against cAMP hydrolysis in the bulk cytosol or at the sarcolemma, we utilized the cAMP FRET biosensors Exchange protein directly activated by cAMP (Epac)-S^H187^ (H187)^22^ and plasma membrane-targeting Epac-S^H187^ (pm-H187). In these sensors, cAMP is detected by the cyclic nucleotide-binding domain within a catalytically dead Epac. Increase in [cAMP] induces a loss of FRET ratio through a conformational change that brings the donor fluorophore CFP away from the two tandem acceptor YFP fluorophores.

In guinea pig cardiomyocytes, the H187 sensor showed a characteristic bulk cytosolic expression with some perinuclear retention (Fig 2A) as reported.^22^ The average baseline ratio of CFP/YFP for the H187 sensor was 0.47±0.0046 (mean±SEM; n=646 cells, N=14 animals; Supplement Fig 2A). The maximum FRET ratio response to the saturating amount of the adenylyl cyclase activator fsk and the pan-PDE inhibitor IBMX was 1.351±1.848. This level was slightly lower than the reported ratio of 1.63.9±10.4,^22^ performed in N1E-115 mouse neuroblastoma cells. The baseline and maximum FRET ratios for H187 (p=0.84 and 0.39, respectively) were not different in male vs female cells. As we have already demonstrated in the mouse,^28^ pm-H187 showed notable membrane localization in guinea pig cardiomyocytes (Fig 2A). The plasma membrane (Supplement Fig 2B) baseline FRET ratio was greater compared to H187 (0.53±0.0028, p<10^-12^, two-tailed Student’s t-test), indicating higher subsarcolemmal [cAMP] at baseline; the maximum FRET change was comparatively reduced (1.024±0.0096, p<10^-12^, two-tailed Student’s t-test). Female cells had lower pm-H187 baseline cAMP level and a greater maximal ratio compared to male counterparts (Supplement Fig 2C).

**Fig 2.**
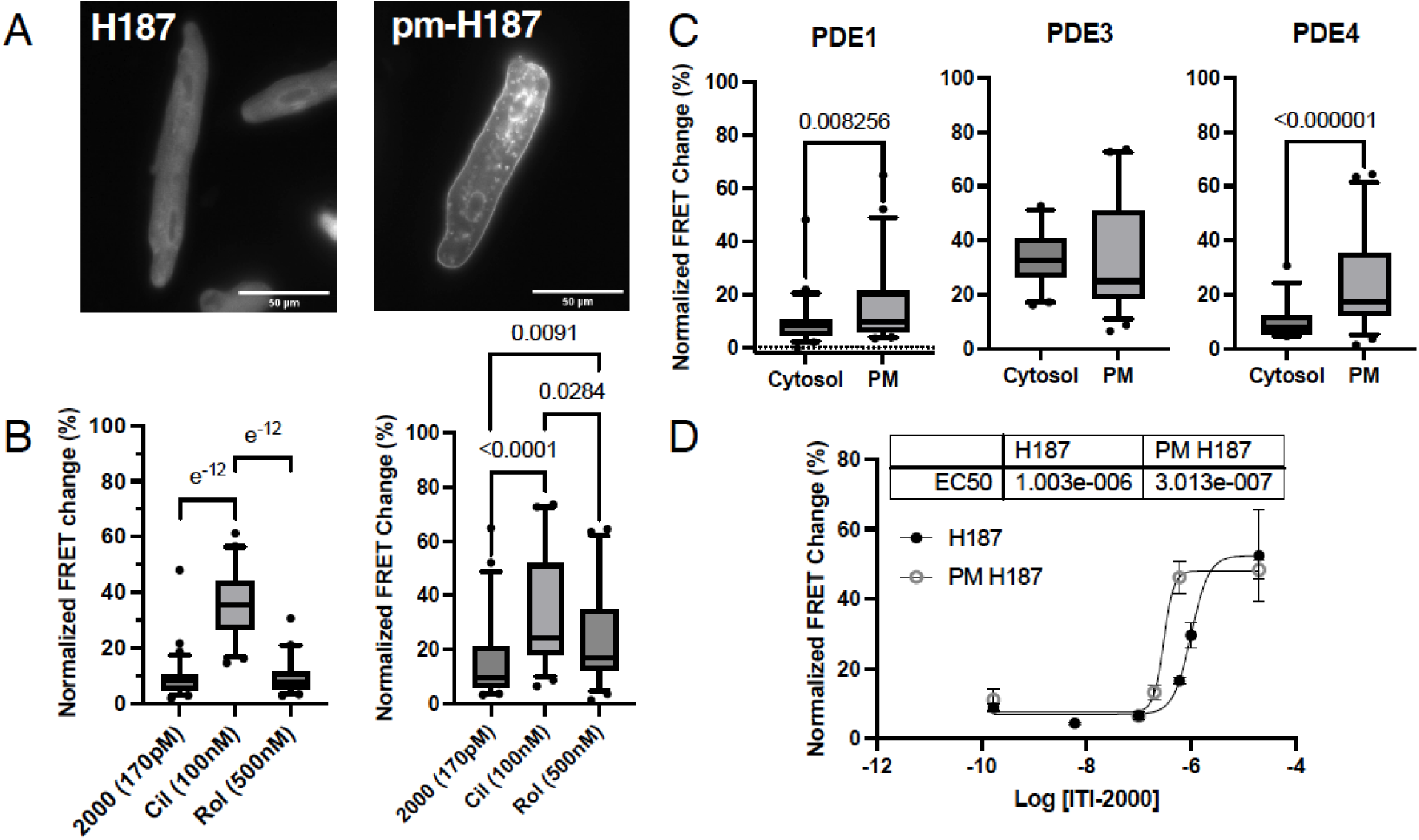
**PDE1 activity at the plasma membrane is greater compared to the cytosol**. A) Micrographs of adult guinea pig cardiomyocytes expressing the cytosolic (H187) or plasma membrane cAMP biosensor (pm-H187). B) Box-whisker plots (5-95^th^ percentile) showing FRET change in response to 2x IC_50_ of PDE1, 3, or 4 inhibitors (respectively, IC200041 (2000), Cilostamide (Cil), or Rolipram (Rol)) using the indicated cAMP biosensor. Kruskal-Wallis test values with Dunn’s analysis shown (H187: n=167, N= 4; pm-H187: n=143, N=8). C) A direct comparison of individual PDEs inhibitors at the cytosol vs plasma membrane. T-test values shown. D) A dose response curve of PDE1 inhibitor as detected by H187 or pm-H187 (n=7-110, N=9). A four parameters least squares fit was performed, and EC_50_ values are shown.

The hydrolytic activities of different PDEs against cAMP were compared at 2x EC_50_ values for PDE1, 3, and 4 specific inhibitors (respectively: IC200041, cilostamide, and rolipram). PDE3 was the most active enzyme, hydrolyzing cAMP at both the cytosol and the membrane (Fig 2B) without a domain-dependent change (Fig 2C, middle). In the cytosol, PDE3 was an especially robust regulator of cAMP in male compared to female cells (Supplemental Fig 2D, left). PDE4D regulates ryanodine receptor at the junctional SR^12^ and indeed showed greater activity at the plasma membrane compared to the cytosol (Fig 2C). PDE1 showed greater regulation of cAMP at the membrane compared to the cytosol in both male and female cells. Likely reflecting the lower level of basal plasma membrane [cAMP] in the female, we noticed diminished responses to all PDE inhibitors in female versus male cells in this domain (Supplement Fig 2D, right). A dose response curve to PDE1i supported this observation by showing a leftward shift in cAMP regulation at the membrane compared to the bulk cytosol (EC_50_ = 300 nM vs 1 µM; p<0.0001, F (1, 375) = 24.50, Fig 2D). The presence of greater cAMP-hydrolyzing PDE1 activity at the membrane (Fig 2C) may reflect what is happening in the bulk cytosol: with greater proportion of the cGMP-hydrolyzing PDE1A found in the cytosol and comparatively less of the cAMP-specific PDE1C (Fig 1B).

PDE1i induces positive inotropic effects in rabbit and guinea pig cardiomyocytes, only in the presence of sub-maximal fsk stimulation of adenylyl cyclase to elevate basal cAMP levels.^5,18^ We replicated these experiments using the H187 cAMP biosensor.

Sub-saturating concentrations of fsk and PDE1i synergistically increased cAMP production more so than the effects of either drug alone (Supplement Fig 2E).

### PDE1 Regulates A_2A_R-Coupled cAMP, and PKA to a Lesser Extent

The inotropic effects of PDE1i in vivo require Ado signaling, and PDE1C multi-complexes with A_2A_R.^29^ We thus tested the hypothesis that PDE1 and A_2A_R regulate the same pool of sarcolemma cAMP. To do so, we performed a series of FRET experiments where we compared the magnitude of change in [cAMP] upon PDE1i in the presence or absence of an A_2A_R ligand or agonist. If an increase in cAMP upon treatment with A_2A_R stimulation would potentiate the effects of PDE1i, then we would consider that as evidence of coordinated regulation between the two proteins. In an initial set of experiments, cells were treated with the A_2A_R specific agonist CGS21680 (CGS). CGS yielded a transient rise in cAMP at doses greater than 1 µM. The peak change in [cAMP] was dose-dependent, with a steep Hill coefficient (8.345) and an EC_50_ of 8.23 µM (Suppl Fig 3A and B). Using a sub-maximal dose of 1 µM, we observed that A_2A_R stimulation significantly increased the FRET response upon PDE1i in the bulk cytosol (Suppl Fig 4A-C). This synergistic regulation where A_2A_R and PDE1 coordinate the regulation of a common cAMP pool was replicated at the sarcolemma using the pm-H187 biosensor (Fig 3A-B). Next, we confirmed that PDE1’s relationship to A_2A_R is unique and not redundant with the other cAMP-hydrolyzing enzymes PDEs 3 and 4.^3,18^ Neither of the latter two PDEs significantly regulated the pool of cAMP associated with A_2A_R whether at the sarcolemma (Fig 3C-F) or at the bulk cytosol (Suppl Fig 4D-G).

**Fig 3.**
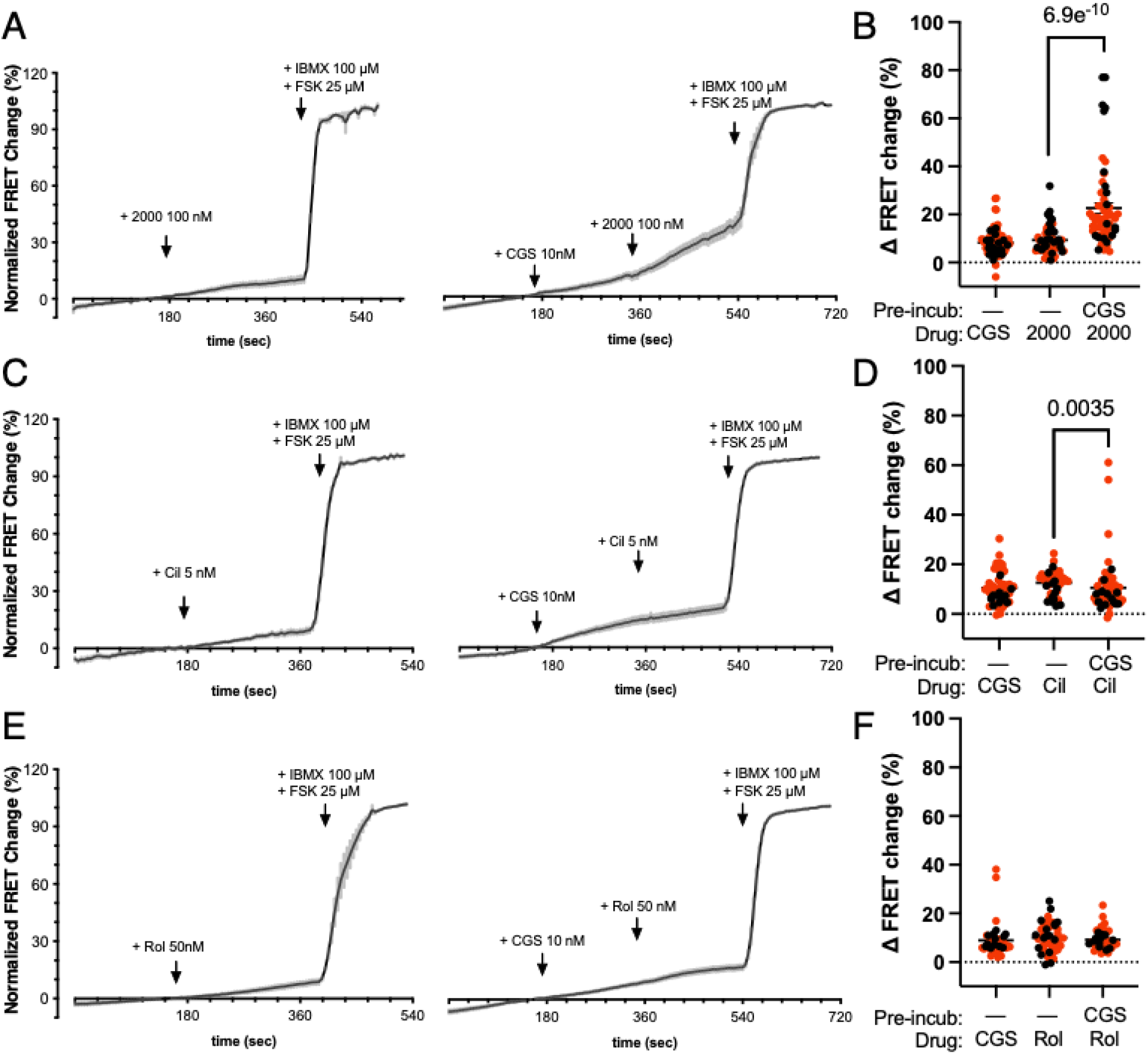
Among cAMP-hydrolyzing PDEs 1, 3, and 4, only PDE1 regulates A_2A_R-stimulated cAMP pool. Healthy guinea pig ventricular cardiomyocytes overexpressing pm-H187 cAMP FRET biosensor were treated with the indicated pharmacological agents. A) Representative FRET tracing of cells expressing pm-H187 cAMP biosensor and treated with PDE1 inhibitor (2000, 100nM) or treated sequentially with CGS (10nM) and 2000 (100nM) and B) corresponding group average results are plotted (n=41, N=9). Males are represented by black dots and females by red dots. C) Representative FRET tracing of cells treated with PDE3 inhibitor (Cil, 5nM) or sequentially with CGS (10nM) and Cil (5nM) and D) corresponding group average results are plotted (n=54, N=4). E) Representative FRET tracing of cells treated with PDE4 inhibitor (Rol, 50nM) or sequentially with CGS (10nM) and Rol (50nM) and F) corresponding group average results are plotted (n=40, N=4). Kruskal-Wallis H tests were significant: B) H(3)=176.6, p<1e-6. D) H(2)=39.07, p<1e-6. F) H(2)=77.77, p<1e-4. Dunn’s post-hoc test p values are shown.

Lower drug doses were required to study A_2A_R/PDE1 signaling at the plasma membrane as not to saturate [cAMP] or overwhelm its local gradient.^7^ At these doses of CGS (100 nM at the membrane and 1 µM at the cytosol), we observed that female cells showed greater drug response at the membrane compared to the cytosol (Suppl Fig 4H).

PKA is the primary effector of cAMP that increases cell contractility and is necessary in the positive inotropic effects of fsk/PDE1i.^5^ We next determined that cAMP regulation by A_2A_R and PDE1 is sufficient to activate PKA as measured by its cytosolic FRET biosensor AKAR4.^24^ Testing a range of 2000 dosage, we saw a dose-dependent PKA activation that showed a large variance ≥ 400nM (Fig 4A). No matter the dose, PDE1i did not further increase A_2A_R-mediated PKA activation (Fig 4B). This was again demonstrated in PKA phospho-substrate (pPKA) staining of fixed cells incubated with PDE1i and/or CGS for five minutes (Fig 4C-D). PDE1i or CGS individually increased pPKA intensity, but did not synergistically increase it. We next hypothesized that PKA activation downstream of A_2A_R/PDE1 may phosphorylate specific contractile or Ca^2+^handling proteins (Suppl Fig 5A and B). All drug treatment conditions failed to induce phosphorylation at any of these sites. Overall PKA activity in cell lysates was also assessed for pPKA intensity (Fig 4E). While the broad β-adrenergic receptor agonist ISO and Fsk+IBMX induced increases in PKA phosphorylation, we did not detect a bulk, sustained increases in phosphorylation for CGS+PDE1i.

**Fig 4.**
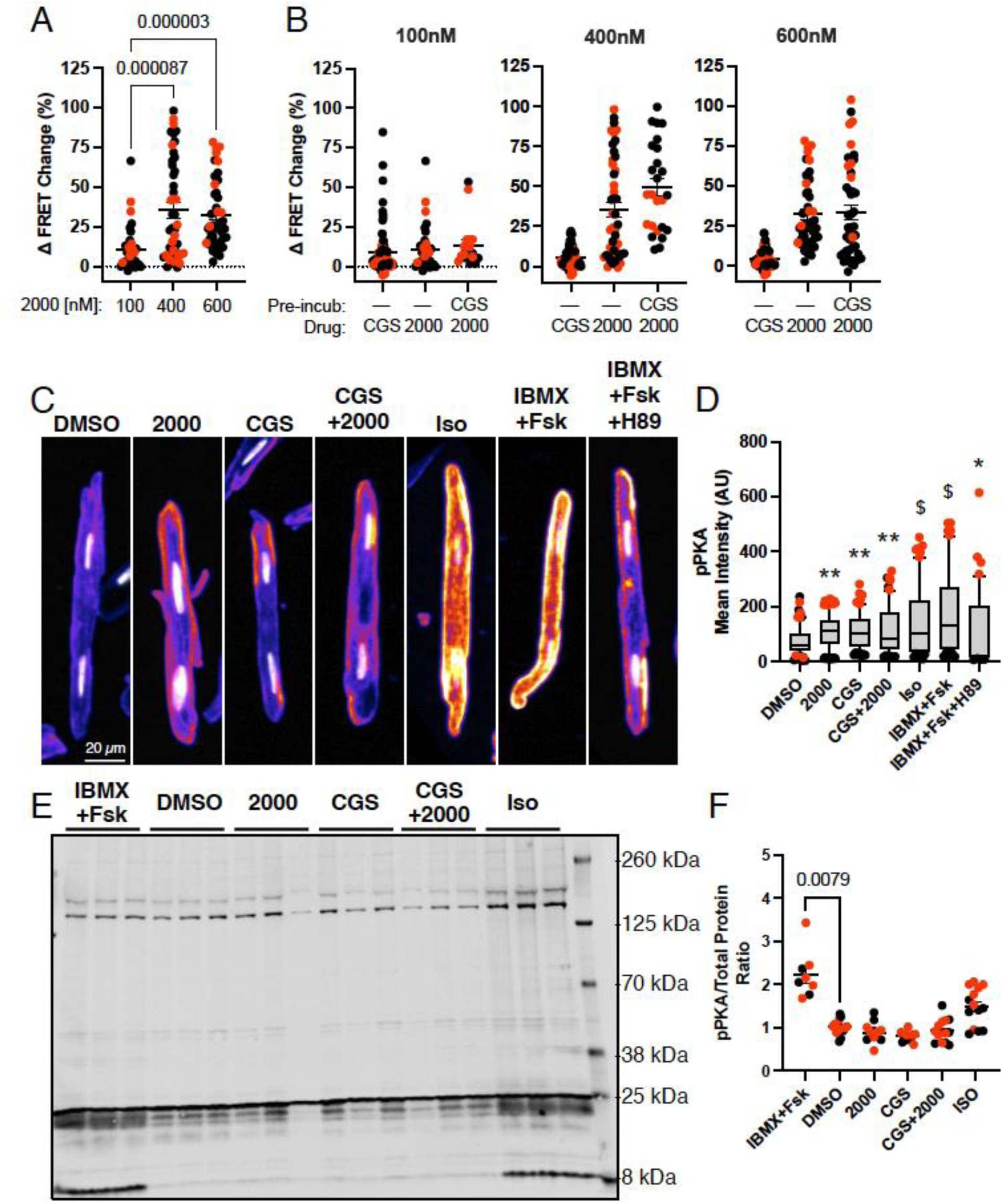
Minimal effects of A_2A_R/PDE1 on PKA activity. Healthy guinea pig ventricular cardiomyocytes overexpressing the AKAR4 PKA activity FRET biosensor were treated with the indicated pharmacological compounds. A) Grouped average FRET response to PDE1i 2000 at 100nM, 400nM, or 600nM (Kruskal-Wallis test, H(2)=26.791, p<2e^-6^, n=33-46 N=15). B) Grouped average FRET response of CGS (1uM), 2000, or sequential treatment with CGS (1uM) and 2000 at different concentrations. Kruskal-Wallis tests were for 100nM: H(2)=6.306, p=0.0427, n=18 N=6; 400nM H(2)=59.09, p=1e^-12^, n=26 N=9;(600nM) H(2)=62.79, p=1e^-12^, n=46 N=9. However, Dunn’s post-hoc test found no significance between the last two bars in each respective graphs. C) Representative micrographs of guinea pig cardiomyocytes treated with respective drugs for 5 minutes, fixed, and stained with PKA phosphorylation (pPKA) substrate antibody. Isoproterenol (Iso) and IBMX+Fsk were used as a positive control, DMSO and H89 were used as negative controls. D) Graph plot for pPKA fluorescence mean intensity of guinea pig cardiomyocytes treated with different drugs. Sidak’s test F(6, 996)=15.58, n=30-60 N=3, and p values are indicated (*p<1^e-6^, **p<0.005, $p<5e^-9^) E) Western blot analysis of pPKA substrates expression in adult guinea pig cardiomyocytes treated with respective drugs. F) Quantification of pPKA western blot analysis plot. (Males are shown in black dots and females are in red dots). Kruskal-Wallis tests were significant: H(5)=35.89 p<0.001.

### Positive Inotropic Effects of PDE1i and A_2A_R Stimulation

Provided that the cAMP/PKA cell signaling pathway is a classic modulator of myocyte contractility, we tested if PDE1 and A_2A_R would exert positive inotropic changes in paced cardiomyocytes. Freshly isolated guinea pig ventricular cardiomyocytes were loaded with the Ca^2+^ sensor Fura-2AM, and cell sarcomere shortening (SS) and intracellular Ca^2+^ transients (CaTs) were monitored using IonOptix.^5,18^ Representative traces are shown in Figure 5A-C. SS or CaT (top or bottom, respectively) traces are shown at baseline or upon exposure to A_2A_R agonist CGS (1 µM) or PDE1 inhibitor 2000 (1 µM) or in combination. In combination, the drugs induced positive inotropic changes without a parallel increase in peak CaT in male and female cells (Fig 5D and E, left). A trend towards or significantly improved lusitropic effect was observed in male and female cells, respectively (Supplement Table 1; p=0.07 and p=10^-7^, respectively, 2-way ANOVA with uncorrected Fisher’s LSD). CGS had minor effects on cell contractility or CaT, no matter the sex. Male cardiomyocytes did not respond to PDE1i, as reported before.^18^ However, 2000 alone increased cell contractility (Fig 5D, right, p=0.00012 vs baseline, p=0.00157 vs male; 2-way ANOVA with uncorrected Fisher’s LSD) and improved relaxation (Supplement Table 1, p=6.1×10^-6^, 2-way ANOVA with uncorrected Fisher’s LSD) in female cardiomyocytes. Peak CaT level was increased in these cells (Fig 5E, right, p=0.012). Even at a lower dose of 2000 (100 nM), PDE1i increased CaT changes to induce positive inotropic effects in female cardiomyocytes (Supplement Fig 6A). CGS did not further augment these effects (Supplement Fig 6B).

**Fig 5.**
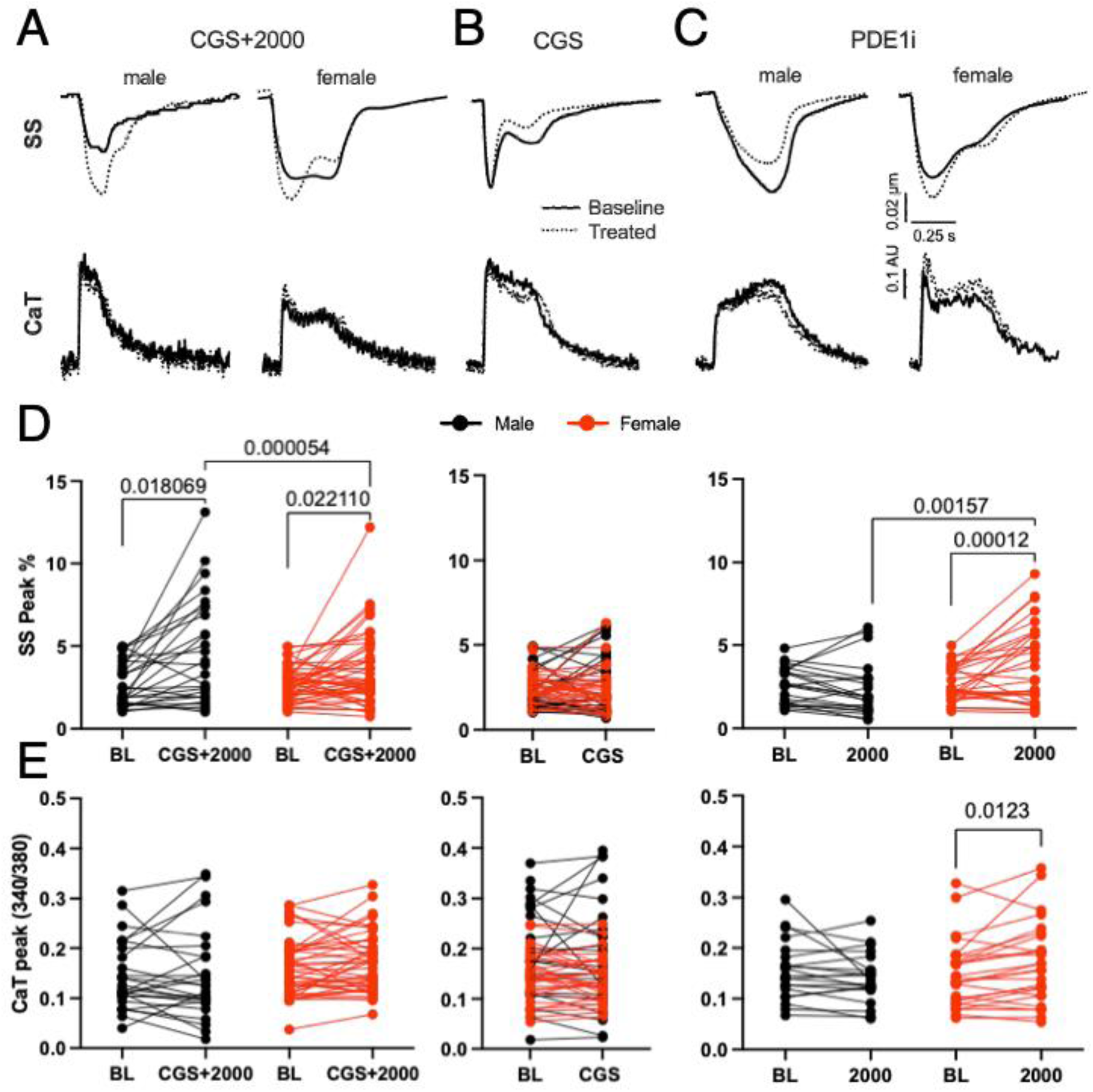
**A_2A_R Stimulation and PDE1i Induce Cardiac Contractility**. Fura-2AM loaded healthy ventricular cardiomyocytes from male and female guinea pigs were electrically paced. Corresponding example traces in cell sarcomere shortening (SS) and intracellular calcium transients (CaT) are shown for each drug treatment. Baseline-treatment responses in D) peak SS % change [(Left: F (1, 73) = 4.111, p=0.046245) and (Right: F (1, 50) = 10.68, p=0.00196), n=27-48 N=14, RM two-way ANOVA, uncorrected Fisher’s LSD post-hoc test p values shown]. The same in E) CaT ratio (340/380) are plotted (Right: F (1, 45) = 8.512; P=0.0055, n=27-48 N=14, RM two-way ANOVA, uncorrected Fisher’s LSD post-hoc test p values are shown).

We further explored the rationale for the sex difference by examining the contribution of the inhibitory G protein A_1_R. Cardiomyocytes were pre-incubated with the A_1_R antagonist DPCPX, and the response to PDE1i recorded. Both the cytosolic and plasma membrane cAMP FRET biosensor revealed a sex-dependent difference in the role of A_1_R to modulate [cAMP] downstream of PDE1i (Supplement Fig 6C and D). In male cells, inhibiting A_1_R led to a greater [cAMP] increase upon PDE1i. This was significantly blunted in female cells (p=1.4×10^-6^ in H187; p=0.0032 in pmH187; 2-way ANOVA with uncorrected Fisher’s LSD).

### PDE1 and A_2A_R Regulate Myocardial Contractility Ex Vivo

We then sought to establish the functional ramifications of the A_2A_R /PDE1 cell signaling pathway at the organ level. The cardiac hemodynamic response to A_2A_R stimulation with CGS (300 nM) alone or upon treatment with PDE1i with 2000 (1 nM) was determined in isolated guinea pig hearts continuously paced at 275 BPM (Fig 6A and Supplement Table 2). These results demonstrate that sequential treatment with PDE1i and A_2A_R agonist significant increased LVDP, contractility (dP/dT_max_), and relaxation (dP/dT_min_). Increas in coronary flow (CF) was primarily influenced by CGS (Figure 6C), thus no significant differences were observed between CGS alone and CGS with 2000. Figure 6D and E demonstrate that the LVDP response to the combined 2000 and CGS was significantly greater than the response to each agent individually (p=0.0112, Kruskal-Wallis test). This observation persisted for both contractility (Fig 6F and G; p=0.0012, 2000 vs 2000+CGS; p=0.0141, CGS vs 2000+CGS, Kruskal-Wallis test), and relaxation (Fig 6H and I; p=0.0040, 2000 vs 2000+CGS; p=0.0016, CGS vs 2000+CGS, Kruskal-Wallis test). Together, these results signified that a synergistic relationship between PDE1 and A_2A_R increases cardiac contractility and relaxation at the organ level.

**Fig 6.**
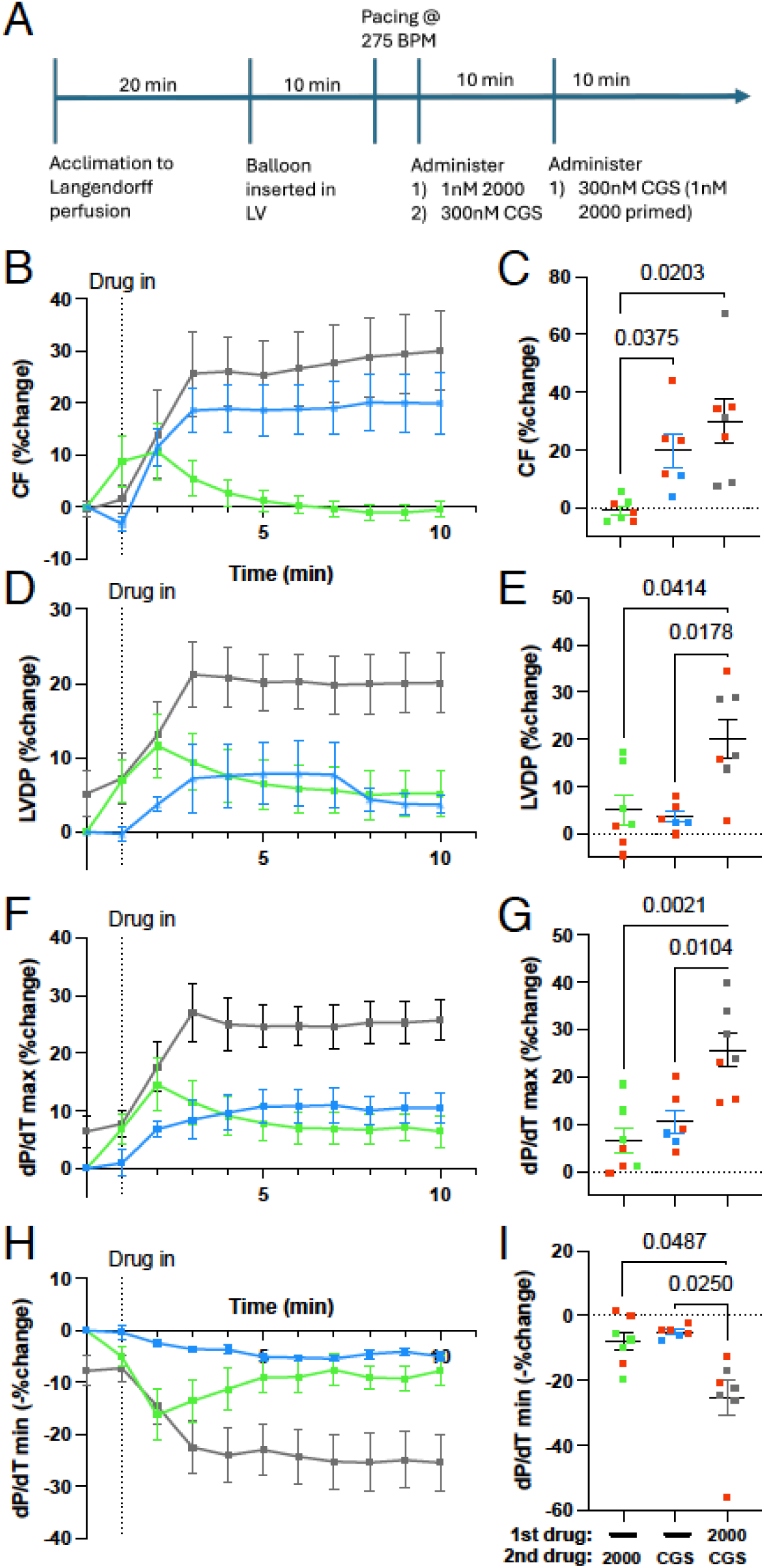
A_2A_R stimulation and PDE1i synergistically increase hemodynamic response in healthy, ex vivo perfused guinea pig hearts. Isolated guinea pig hearts with a balloon pressure catheter inserted into the left ventricle (LV) were treated with the indicated pharmacological agents. A) Timeline of cardiac preparation and treatment. B) Coronary flow (% change) after treatment with 2000 (1nM; green), CGS (300nM; blue), and CGS (300nM) after ten minutes of priming with 2000 (1nM; grey). C) Corresponding group averages at the 10 minute mark are plotted (Brown-Forsythe ANOVA F(2, 11.07) = 8.219, p=0.0065). D) LV developed pressure (LVDP) across the three treatment groups, and E) corresponding group averages are shown (Brown-Forsythe ANOVA F(2, 12.20) = 8.786, p=0.0043). F) The maximum rate of pressure rise (dP/dT_max_) change across time, and G) corresponding group averages are plotted (Brown-Forsythe ANOVA F(2, 15.68) = 12.66, p=0.0005). H) Change in the minimum rate of pressure change (dP/dT_min_) across time, and I) corresponding group averages are plotted (Brown-Forsythe ANOVA F(2, 9.375) = 9.727, p=0.0052). Dunnett’s T3 post-hoc test p values are shown (N=6-7).

### PDE1 and A_2B_R Do Not Regulate a Common cAMP Pool or Cell Contractility

At high concentrations, Ado can also bind to A_2B_R to stimulate cAMP production via G_s_ signaling and exert a mild inotropic response.^30^ This led us to speculate the regulatory role of PDE1 over cAMP pool generated by A_2B_R. Similar to CGS, the A_2B_R-specific agonist BAY60-6583 (BAY) in guinea pig cardiomyocytes induced a transient increase in cAMP at doses greater than 1 µM (Supplement Fig 7A). Plotting the peak response yielded a dose response curve with an EC_50_ of 1.06 µM (Supplement Fig 7B).

Subsequent FRET experiments were performed using 100 nM. At this dose, A_2B_R stimulation did not alter the effects of PDE1i (Fig 7A and B). Isolated cardiomyocytes were again paced to study the functional ramifications of this signaling pathway. Male and female cells were not different in their lack of response to BAY alone or in combination with 2000 (Fig 7C-E and Supplement Table 3).

**Fig 7.**
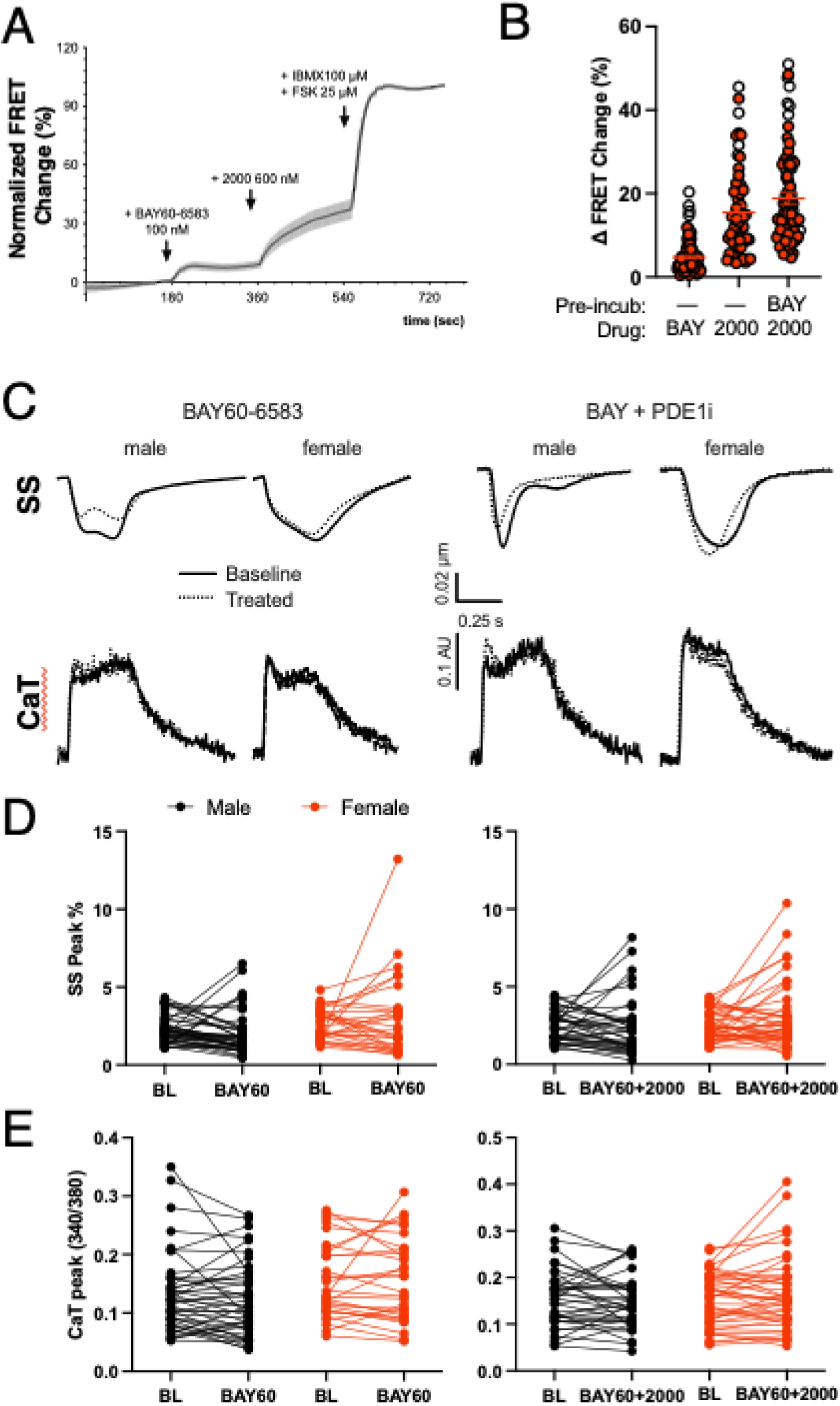
**PDE1 does not regulate A_2B_R-stimulated cAMP pool to induce contractile effects in ventricular myocytes**. Healthy guinea pig ventricular cardiomyocytes overexpressing cytosolic H187 cAMP FRET biosensor were treated with indicated pharmacologic agents. Representative FRET tracing of ventricular cardiomyocytes treated A) sequentially with A_2B_R agonist BAY60-6583 (BAY60, 100nM) and 2000 (600nM). B) Corresponding group averaged results. Kruskal-Wallis test was not significant. C) Example traces for cell SS (top) and intracellular CaTs (bottom) are shown for indicated drug treatment in the male and female guinea pig ventricular cardiomyocytes. Baseline-treatment responses in peak D) SS and E) CaT are plotted. RM two-way ANOVA tests showed no significant interaction between drug response and sex (n=30-45 cells from N=9).

### Functional Coupling Between PDE1 and A_2A_R in HF

PDE1i increases myocardial contractility in patients with HFrEF.^6^ We tested the ramifications of our functional findings in a guinea pig model of HFrEF. Only males were included owing to heterogenous surgery response in female animals.^31^ In this model adapted from Liu et al.^19^ and described in the Methods, ascending aortic constriction (AC) increases the afterload on the LV. At study termination, the AC group had reduced LV function as assessed by EF and FS compared to the Sham group, without differences in the body weight or HR. Structurally, the group had significantly greater LV weight as adjusted for the body weight (p=0.01, Mann-Whitney) and had a trend towards or significant chamber dilation and wall thinning (Supplement Table 4). In cardiomyocytes isolated from the Sham group, PDE1i induced lusitropic effects in SS along with enhanced Ca^2+^ reuptake rate (supplement Table 5). A_2A_R stimulation had a mild Ca^2+^-independent positive inotropic effect. When combined, Ca^2+^-independent inotropic and improved contractile kinetics but no lusitropic effects were observed (Supplement Table 5). In cells isolated from guinea pigs with HF, Ca^2+^-driven positive inotropic and lusitropic effects were evident only with both PDE1i and A_2A_R stimulation, indicative of a preserved synergistic relationship between the two proteins in HF (Fig 8A and D). As shown in Fig 8B and C, neither PDE1i nor A_2A_R stimulation alone induced positive inotropic or lusitropic changes. On the other hand, A_2B_R stimulation – whether alone or in combination with PDE1i – did not alter cell contractility in failing cardiomyocytes (Supplement Table 6). In contrast, no functional effects were observed in Sham cells treated with A_2B_R stimulation alone or in combination with PDE1i (Supplement Table 5).

**Fig 8.**
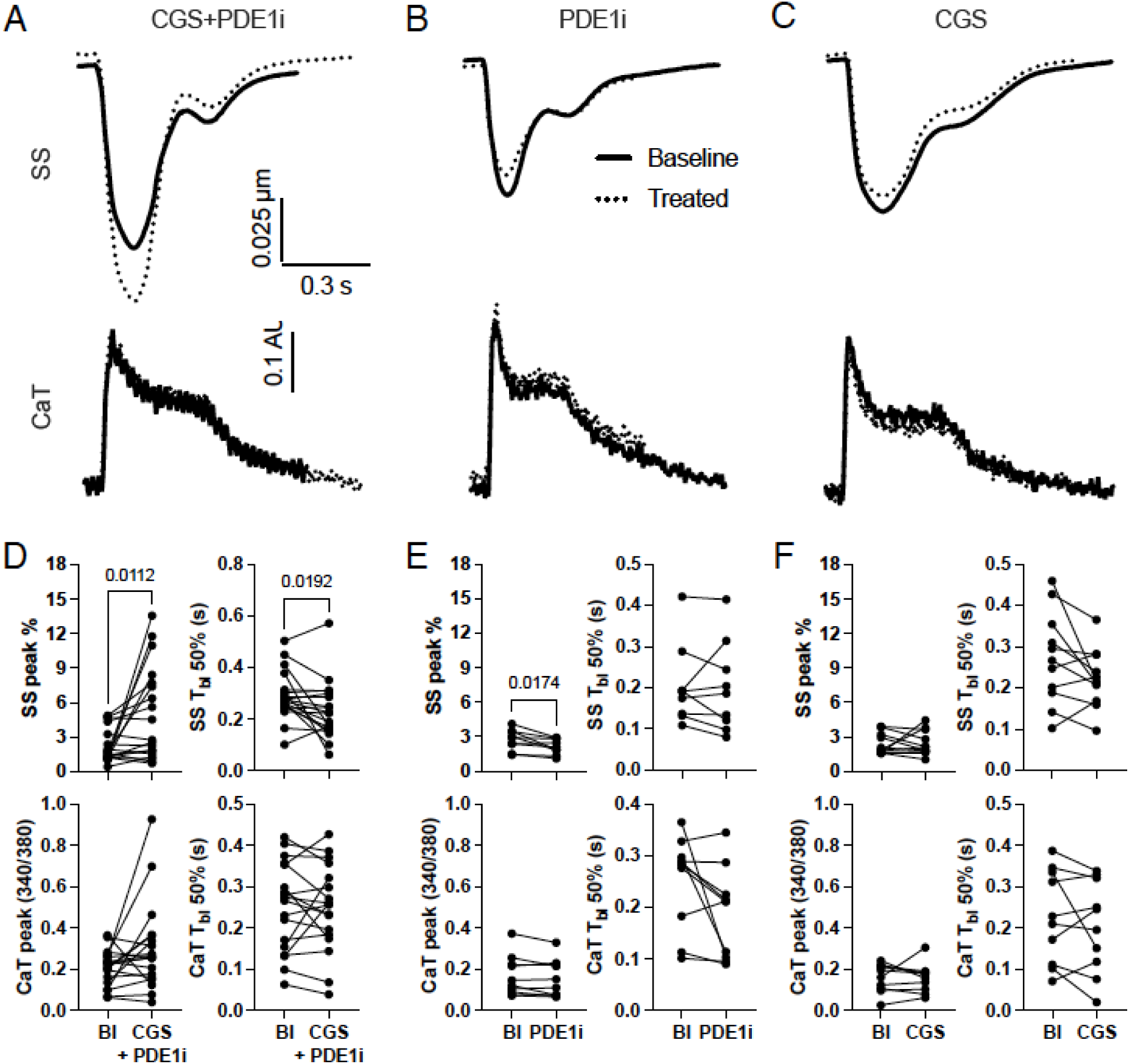
**Contractile effects of A_2A_R and PDE1 modulation are preserved in HF**. Ventricular cardiomyocytes were isolated from male guinea pigs with HF. Sarcomere shortening (SS) and intracellular Ca^2+^ transients (CaTs) were recorded in paced cells. Baseline-treatment responses to CGS+2000 are shown for A) SS and B) CaT parameters. Two-tailed paired T-test p values are indicated (n=30-32, N=5). C) SS and D) CaT peak changes are likewise plotted in response to BAY+2000. Two-tailed paired T-test p values are indicated (n=10-16, N=5)

## Discussion

An interaction between PDE1C and A_2A_R has been observed, in that the two proteins form a complex together to induce cardioprotective effects in mice.^29^ However, PDE1C expression is low in mice,^5^ in which PDEs 3 and 4 are instead predominant regulators of myocardial cAMP.^3^ PDE1i does not affect contractile functions in mice,^5^ whereas it acutely induces inodilatory effects to improve heart performance in human patients with HFrEF.^6^ In human and guinea pig myocardium, PDEs 1 and 3 indeed account for 82-94% of total cAMP hydrolysis.^3^ How PDE1C and A_2A_R regulate cell signaling changes to affect myocardial contractility thus remained unknown in a relevant preclinical model.

Our current findings reveal a presence and role of PDE1C in the sarcolemma membrane that controls a specific pool of cAMP associated with A_2A_R in the guinea pig. A_2A_R-associated cAMP is specifically hydrolyzed by PDE1 and not by PDE 3 or 4. PDE1 does not hydrolyze A_2B_R-associated cAMP. Functionally, the cAMP signaling pathway regulation by A_2A_R and PDE1 has inotropic and lusitropic effects at the cellular and organ levels, and persist in a state of pressure overload HF.

### Evidence of PDE1 localization and activity at the sarcolemma

PDE1C staining was diffuse with pronounced localization at the Z-line in guinea pig cardiomyocytes. An earlier study in the human myocardium, in contrast, reported PDE1C staining of both Z-and M-line structures.^4^ These differences may be due to species or antibody differences, although in our studies we used a commercially available antibody recognizing the C-terminus that is common to all PDE1C splice variants. Additionally, we observed intercalated disc staining of PDE1C. This group of PDE1C was likely immunoblotted in the Cav-3 enriched, microsomal fraction of the myocardium. FRET experiments further corroborated stronger regulation of cAMP at the membrane vs the cytosol in individual cells upon PDE1i. This is unexpected from the finding that PDE1 activity was negligible in the microsomal fraction.^4^ However, together with our findings, the emerging finding is that PDE1, and in particular PDE1C, may be at the plasma membrane. These lines of evidence include: PDE1C complexes with the membrane proteins A_2A_R and TRPC3,^29^ localizes to leading-edge protrusions in polarized human arterial smooth muscle cells,^32^ regulates Ca_v_1.2 activity in a PKA-dependent manner,^18^ and PDE1C is biotinylated within proximity of Ca_v_1.2.^33^ Interestingly, we also observed a nuclear fraction of PDE1C. The role of PDE1C at the nucleus remains understudied, though there is evidence of other cAMP hydrolyzing enzymes like PDE3A2 mediating anti-hypertrophic cell growth response.^13^

PDE1A was predominantly found in the soluble and to a lesser extent in the microsomal fraction of the myocardium. PDE1A is a cGMP-hydrolyzing enzyme and may indeed be the predominant regulator of cytosolic cGMP found in human myocardium.^4^ The preponderance in the soluble vs the microsomal fractions is also consistent with the observation that PDE1A regulates the cGMP pool associated with soluble guanylyl cyclase but not the particulate form in the mouse cardiomyocyte.^34^ At the nucleus, PDE1A regulates the cell cycle and apoptosis to control growth and survival in vascular smooth muscle cells.^35^ Such role for PDE1A is consistent with our observation of the protein in the nuclear fraction.

### PDE1C and A_2A_R are functionally coupled at the membrane

GPCRs and PDEs coordinate the regulation of a specific pool of cAMP.^7^ PDE1 does not coordinate β-adrenergic pool of cAMP.^5,6,18^ In isolated guinea pig myocytes, fsk stimulation was necessary to unmask the inotropic effects of PDE1i. These effects were seen within seconds, suggesting that a certain pool of cAMP was readily available to and relevant in PDE1 cell signaling. Our imaging studies provide direct evidence that PDE1 regulates the sarcolemmal cAMP pool generated by A_2A_R but not A_2B_R stimulation in individual cells. These findings are noteworthy in that we provide a) temporal resolution of cAMP in b) different domains within c) a cell whose cAMP signaling landscape matches that of the human myocardium. Furthermore, we demonstrate the acute contractile ramifications of this signaling pathway, which may be temporally distinct from the role of extracellular cAMP signaling to confer cardioprotection against oxidative stress.^15^ We had previously shown that antagonizing A_2B_R (MRS1754) abrogates the inotropic effects of PDE1i in the intact rabbit heart.^5^ A_2A_R was not blocked in that study, but our current evaluation of both receptors suggest that A_2B_R stimulation is insufficient to induce contractile changes in combination with PDE1i. Possible explanations include non-specific targeting of A_2A_R by MRS1754, or modulation of the drug response by circulating Ado in the intact animal.^5^

### Potential sex-dependent differences in A_1_R, A_2A_R, and PDE1 signaling

We observed that PDE1 regulation of cAMP can leverage myocardium contractile and relaxation function. However, in male compared to female isolated cells, different conditions were needed to observe this effect. In female cells, PDE1i alone was sufficient, with no further effects observed from A_2A_R stimulation. However, A_2A_R stimulation was required in males. Corollary FRET imaging experiments did not show sex-dependent differences in cAMP or PKA changes. Nevertheless, female cells had a lower cAMP level and greater maximal capacity at the sarcolemma compared to male cells, suggestive of a baseline difference in cAMP landscape. Female cells were paradoxically less sensitive to IC200041 vs male counterparts in these experiments at a low drug dose (170 pM), but this may reflect a lower abundance of sarcolemmal cAMP at baseline that is available as a substrate for the enzymatic activity. Alternative interpretations are that activities of A_2A_R may be greater at the plasma membrane in the female while that of A_1_R is greater in the male. Indeed, we observed that A_2A_R stimulation induced a greater level of sarcolemmal vs cytosolic cAMP in females.

Blocking A_1_R potentiated the effects of PDE1i in the male selectively. One possible explanation is the role of sex hormones. Estradiol can increase A_1_R and A_2A_R expression levels in an estrogen-positive human breast cancer cell line.^36,37^ Still other factors may be relevant, as sex differences in myocardial PDE4D protein expression heterogeneity^38^ and estrogen regulation of Ca_v_1.2 have been reported.^39^

Still, we did not observe a consistent sex-based difference in freshly prepared ex vivo hearts. To our knowledge, there is no precedence for sex-dependent differences in PDE1 signaling in the healthy myocardium. In our prior studies in dogs, rabbits, mice,^5^ and guinea pigs,^18^ only males were included in the study. Bork et al.^34^ utilized both male and female mice, but no sex-dependent effects were reported. In Gilotra et al.,^6^ female patients made up 42% of the study population (N=27) and responded similarly as male patients did. Interestingly, heterodimer formation between the two ARs have been observed in the striatum of the brain, with reports suggesting that the heterodimers may alter the response to or affinity for drugs that are designed to be specific for one of the two ARs.^40^ Whether A_1_R /A_2A_R heterodimers exist in the myocardium and if PDE1C may regulate the cAMP production associated such heterodimers remain to be demonstrated. We were unable to further determine sex-dependent dimorphism in the HF study, as females have heightened heterogeneous response to AC surgery.^31^

### Mechanism of cell contractile increase upon A_2A_R stimulation

There is conflicting evidence on the inotropic effects of A_2A_R stimulation. In the rat, A_2A_R stimulation increased inotropic effects in cardiomyocytes,^17^ with Ado showing the same effect ex vivo in a manner sensitive to an A_2A_R antagonist.^30^ A mouse model with A_2A_R deletion further confirmed these findings.^41^ However, no inotropic effects were observed in other studies utilizing the rat, guinea pig,^42^ and/or rabbit papillary muscles or cardiomyocytes.^30^ In guinea pig, one study reported increases in cardiomyocyte [cAMP]^42^ while another reported no changes.^43^ These studies, however, evaluated a biochemical snapshot evaluation of [cAMP] in drug-treated cell lysates. In addition to species differences, some of the discrepancies may also derive from the basal adrenergic tone as determined by [cAMP] or load conditions inherently different in intact or isolated myocyte experiments. A_1_R induces anti-adrenergic response in contractility,^14^ and we did not determine A_1_R activity differences in the ex vivo hearts.

Thus, potential differences in [cAMP] or load may explain our observation that intact myocardium treated with A_2A_R agonist showed positive inotropic and lusitropic effects, while no such effects were observed in isolated, unloaded cardiomyocytes. The small increase in cAMP with CGS at 1 µM in our FRET experiments correlated with a minor functional effect in cardiomyocytes. Interestingly, we noted that A_2A_R stimulation induced a transient increase in cAMP at concentrations ≥1 µM that peaked and returned to near baseline level over the course of 3 minutes (Suppl Fig 3E). Such kinetics may represent nonspecific drug responses (possibly activating A1R),^17,41^ or receptor desensitization.^30^

Where an A_2A_R-induced inotropic response was previously observed, Ca^2+^-independent mechanisms have been proposed.^44^ Replicating our prior observation using the nonphysiological agonist fsk,^18^ a parallel change in peak CaT was likewise absent in the current study where A_2A_R stimulation and PDE1i induced inotropic effects in healthy and failing myocytes. In our investigation of the inotropic mechanism in healthy myocytes, we did not find evidence to support PKA activation in individual myocytes as well as in bulk cell lysates. Moreover, we failed to detect drug-induced changes in PKA protein targets such as MyBP-C, TnI, and PLN. These PKA targets were unchanged in guinea pig cardiomyocytes treated with fsk and PDE1i.^5^ It may be that we have yet to elucidate (a) particular PKA target(s) that critically mediates the effects of A_2A_R stimulation and PDE1i. Alternatively, the significance of cAMP as a mediator of A_2A_R-induced contractile changes has been challenged.^17,45^ PDE1 may induce inotropic effects via alternative signaling mechanisms such as Epac,^46^ and warrant further evaluation.

### Clinical Relevance in the Heart and the Brain

Our findings have clinical relevance as the PDE1 inhibitor lenrispodun has completed several clinical trials in patients with HFrEF or Parkinson’s Disease (PD) with a safe profile.^47^ Circulating Ado levels are elevated in patients with chronic HF,^48^ cardiogenic shock,^49^ and ischemic heart disease.^50^ If [Ado] is sufficiently high to stimulate A_2A_R signaling, PDE1i effects may be greater under such pathologic conditions. Ado levels may thus become a relevant factor for consideration in titrating patients with PDE1i.

However, Ado is rapidly hydrolyzed, and its influence may be confined to a narrow window.^14^ Although of a small group of 27, the clinical trial in HFrEF patients included 9 patients with ischemic HF.^6^ The study did not indicate any differential efficacy of PDE1i depending on ischemic vs non-ischemic forms of HF.

PDE1i also induces temporary vasodilatory effects to reduce systemic vascular resistance and lower both systolic and diastolic arterial blood pressure (BP).^6^ A_2A_R agonist is a potent vasodilator of its own, and a combined interaction may potentially induce hypotension and exceed the desired effect of afterload reduction from PDE1i alone. In our study, we observed that at the clinically relevant dose of 1nM, PDE1i only transiently increased the coronary flow in the constant pressure Langendorff perfusion experiments. On average, PDE1i increased the vasodilatory effects of A_2A_R stimulation, though this was not statistically significant (p=0.167). These results together indicate that PDE1 and A_2A_R may not strongly or persistently induce hypotension. As a caveat, HR was kept constant in our study, and the ex vivo study lacked global mechanisms of BP regulation such as baroreceptor reflex. Thus, future studies may be necessary to fully test the vasodilatory effects in vivo and with chronic exposure.

Ado infusions are used for diagnostic purposes. This includes infusion of the A_2A_R agonist regadenoson in pharmacologic stress testing and therapeutically with the administration of rapid IV boluses of Ado for acute termination of a supraventricular tachycardia. A_1_R of nodal cells largely mediate the termination by temporarily blocking reentrant waves via the atrioventricular node.^30^ This observation raises the possibility that in a hypothetical scenario where regadenoson or Ado infusion is given in the background of PDE1i, there may be greater inotropic or yet different effects.

PDE1i is under active investigation for consideration as an adjunctive in PD patients with motor fluctuations and levodopa treatment-induced dyskinesia (NCT05766813). Here, the therapeutic efficacy of PDE1i is most likely attributable to the enzymatic regulation of cGMP as opposed to that of cAMP via the isoform PDE1B.^51^ PDE1B is absent in the heart,^52^ but the enzyme’s regulation of likely nitric oxide and sGC-associated cGMP may represent a unique compartmentalized signaling of the central nervous system. While PDE1C is expressed in the nucleus accumbens to a much lower extent (by a 10-fold) compared to PDE1B,^51^ A_2A_R is considered an important non-dopaminergic regulator of cAMP in the medium spiny neurons (MSNs) of the indirect output pathway.^53^ PDE10A is highly expressed in these indirect MSNs that express A_2A_R and dopamine 2 receptor (D_2_R).^9^ Recently, Chen et al. showed PDE10A regulates cAMP pool associated with A_2A_R-D_2_R heteromerization in the mouse heart.^54^

### Limitations and Conclusion

Our studies are limited in that the PDE1 inhibitor does not discriminate between PDE1A and 1C isoforms. Bork et al. showed that PDE1A regulates cGMP to modulate cell relaxation in the mouse myocyte. We did not assess changes in cGMP levels in our experiments. Although PDE1A has a supraphysiologic K_m_ value of 113 µM for cAMP,^52^ we cannot rule out the contribution of PDE1A as an additional regulator of cAMP other than PDE1C. Our study relied on pharmacologic rather than genetic techniques.

Nevertheless, the finding that PDE1i and A_2A_R but not A_2B_R agonist regulate common pools of cAMP and myocardial contractility support our conclusions. Considerations of sex differences in the setting of HF is being evaluated as part of another study. Taken together, we describe a sarcolemmal domain within which PDE1 and A_2A_R, but not A_2B_R, work to regulate cAMP signaling pathway and control cardiac contractile function. This conclusion is congruous with many prior works summarized in recent reviews^9,55^ that describe specific ways in which cAMP signaling is compartmentalized within the cardiomyocyte.

## ACKNOWLEDGEMENTS

We thank Jake Cunningham for assistance with optimizing myocardial fraction protocol, Dr. James Szczerkowski for assistance with phospho PKA imaging analysis, Alejandro Sanchez for critical reading of the manuscript, and Dr. Sakthivel Sadayappan for generously sharing MyBP-C phospho-specific antibodies.

## SOURCES OF FUNDING

Research reported in this publication was supported by: Achievement Rewards for College Scientists, Inc. (ECC); the National Heart, Lung, And Blood Institute of the National Institutes of Health, under awards R01 HL146169 (MWK), R01HL171586 (GKM), and R01HL171586-02S1 (GKM); and the British Heart Foundation, grants RG/17/6/32944, PG/23/11321 and PG/21/10611 (all to MZ).

## DISCLOSURE

Intra-Cellular Therapies, Inc., provided IC200041 at free-of-charge under an MTA. MLF, MAMG, MT, EDK, ECC, PK, EJ, SS, OS, MZ, MWK, and GKM have nothing to disclose.

## SUPPLEMENTAL MATERIAL

Tables S1–S6

Figure S1-S7

Major Resources Table

---

**Supplemental Figure 1.**
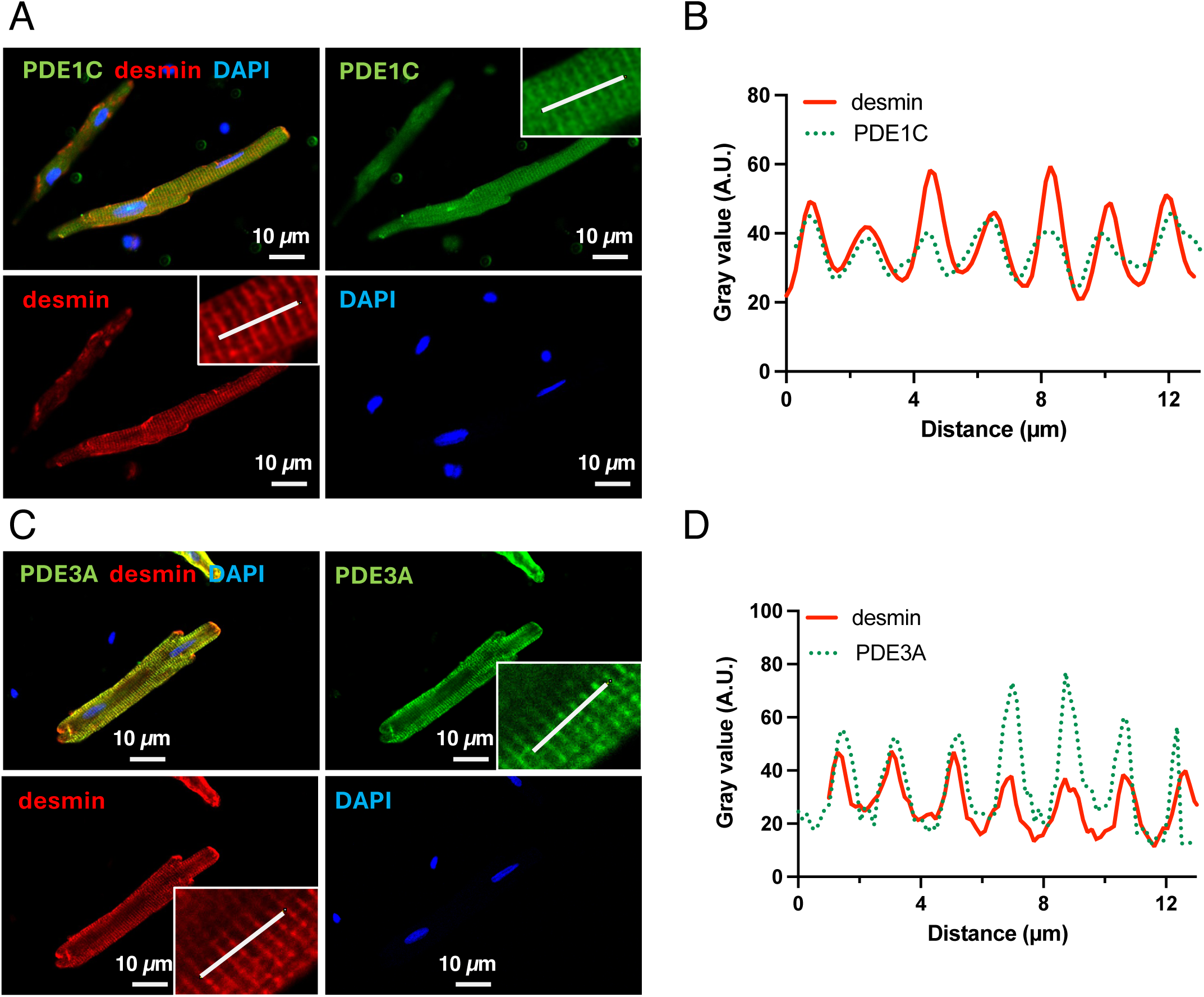
Representative immunocytochemistry staining of A) PDE1C and C) PDE3A in adult guinea pig ventricular cardiomyocytes co-stained with desmin and DAPI. Intensity line plots of desmin and B) PDE1C or D) PDE3A along the line indicated in A and C. Representative out of triplicate experiments.

**Supplemental Figure 2.**
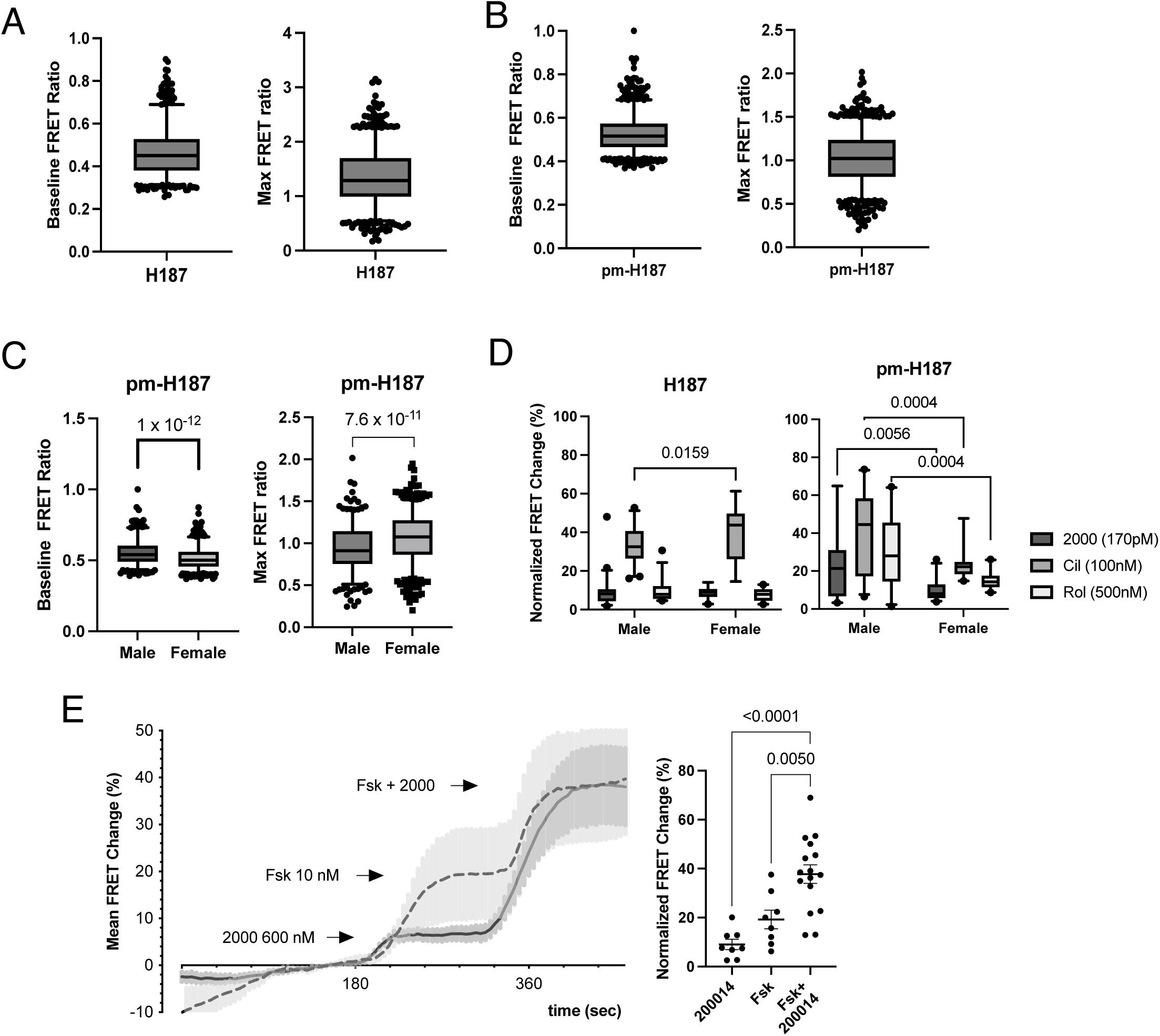
Baseline and maximum FRET ratios in adult guinea pig cardiomyocytes expressing A) the cytosolic cAMP FRET biosensor H187 (n=788, N=14) or B) sarcolemmal pm-H187 (n=923, N=14) shown as box and whisker plots. C) Comparison of baseline and max FRET change in male vs female using pm-H187 sensor. Student’s t-test p values shown. D) Normalized FRET change comparison for 2x IC_50_ inhibitors of PDE1 (2000), PDE3 (cilostamide), or PDE4 (rolipram) in male vs female cells expressing H187 and pm-H187 sensor. Two-way ANOVA: H187: F (2, 165) = 3.382, p=0.0363, pm-H187: F (2, 135) = 0.1876, p=0.8292 with Sidak’s post-hoc test was performed. Significant p values are shown. E) Representative trace showing FRET response to indicated drugs: 2000 or forskolin (Fsk), and group average dot plots shown to the right. Ordinary One-way ANOVA: F (2, 29) = 15.71, p=0.000023. Significant p values are shown following Tukey’s post-hoc test.

**Supplemental Figure 3.**
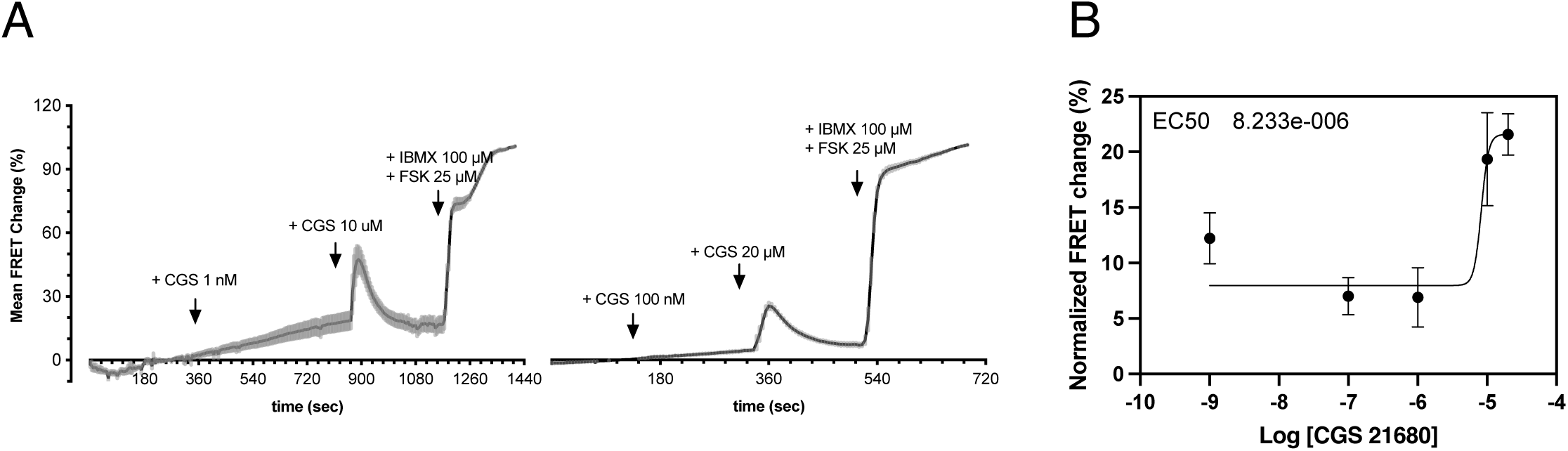
A) Guinea pig cardiomyocytes expressing the cytosolic cAMP sensor H187 were treated with various CGS21860 (CGS) dose. B) Log-transformed dose response curve of CGS 21680 fitted with 4-parameter variable slope least squares fit (n=8-22, N=2).

**Supplemental Figure 4.**
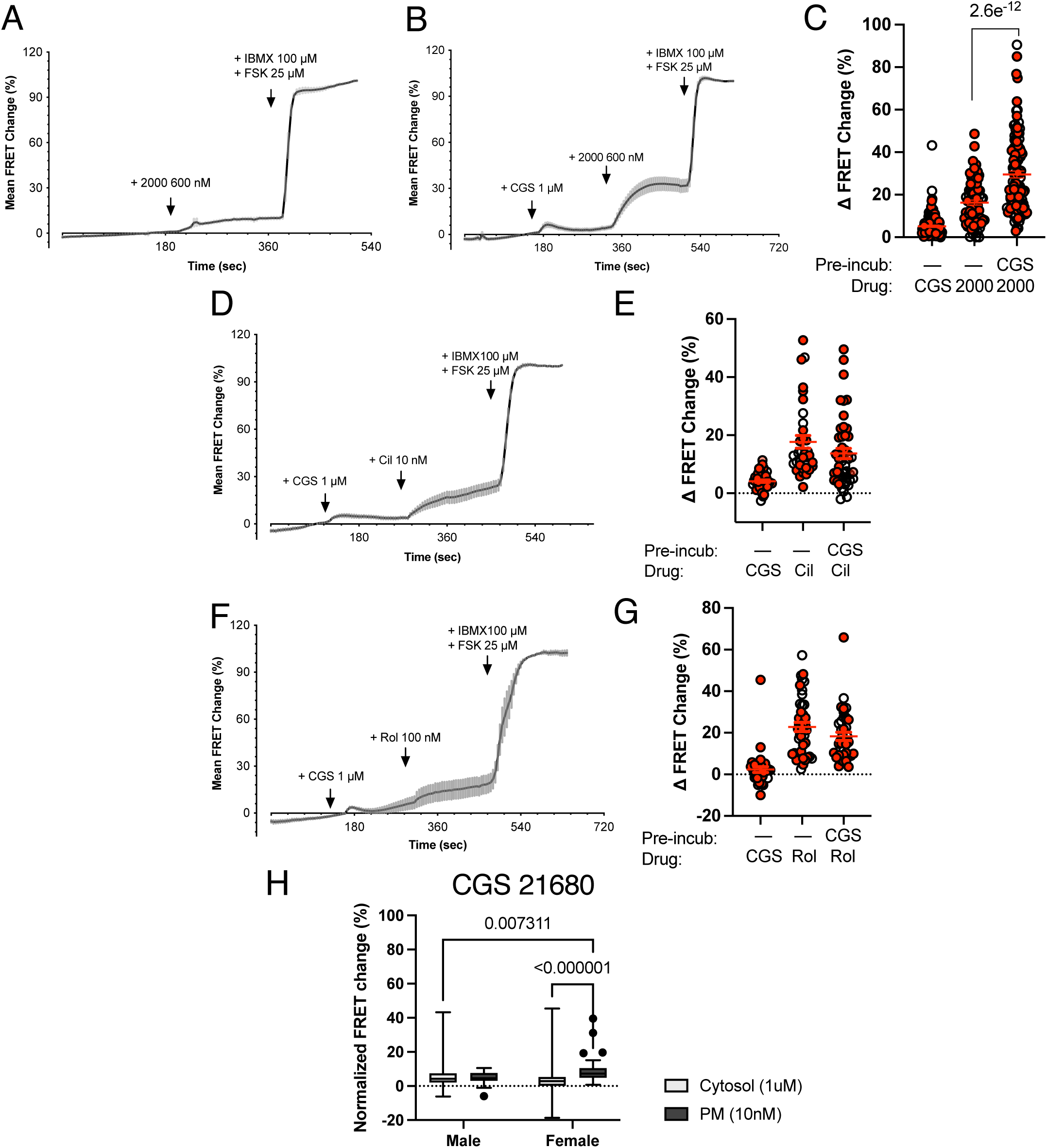
Guinea pig ventricular cardiomyocytes overexpressing H187 cAMP FRET biosensor treated with indicated compounds: A and B) 2000 (600nM) with or without prior CGS (1µM). C) Group averaged FRET response (Ordinary One-way ANOVA, F (2, 314) = 115.33, p<10^-13^, n=78-121, N=4-5, Holm-Sidak’s post-hoc test value shown.) Representative FRET tracing of ventricular cardiomyocytes treated with D) CGS, cilostamide (Cil, 10nM) and E) corresponding group averaged FRET responses (Ordinary 1-way ANOVA, F (2, 125) = 20.238, p<0.00001, n=27-47, N=4, the last two groups were not significantly different Holm-Sidak’s test). Representative FRET tracing of ventricular cardiomyocytes treated with F) CGS, rolipram (Rol, 100nM) and G) corresponding group averaged FRET responses (Ordinary 1-way ANOVA, F (2, 113) = 31.293, p<0.00001, n=35-42, N=4, the last two groups were not significantly different, Holm-Sidak’s test). H) Normalized FRET response to CGS21680 in male vs female cells expressing the cytosolic H187 vs plasma membrane pm-H187. Two-way ANOVA, F (1, 266) = 11.65, p=0.000741 with Tukey’s post-hoc test. Significant p values are shown.

**Supplemental Figure 5.**
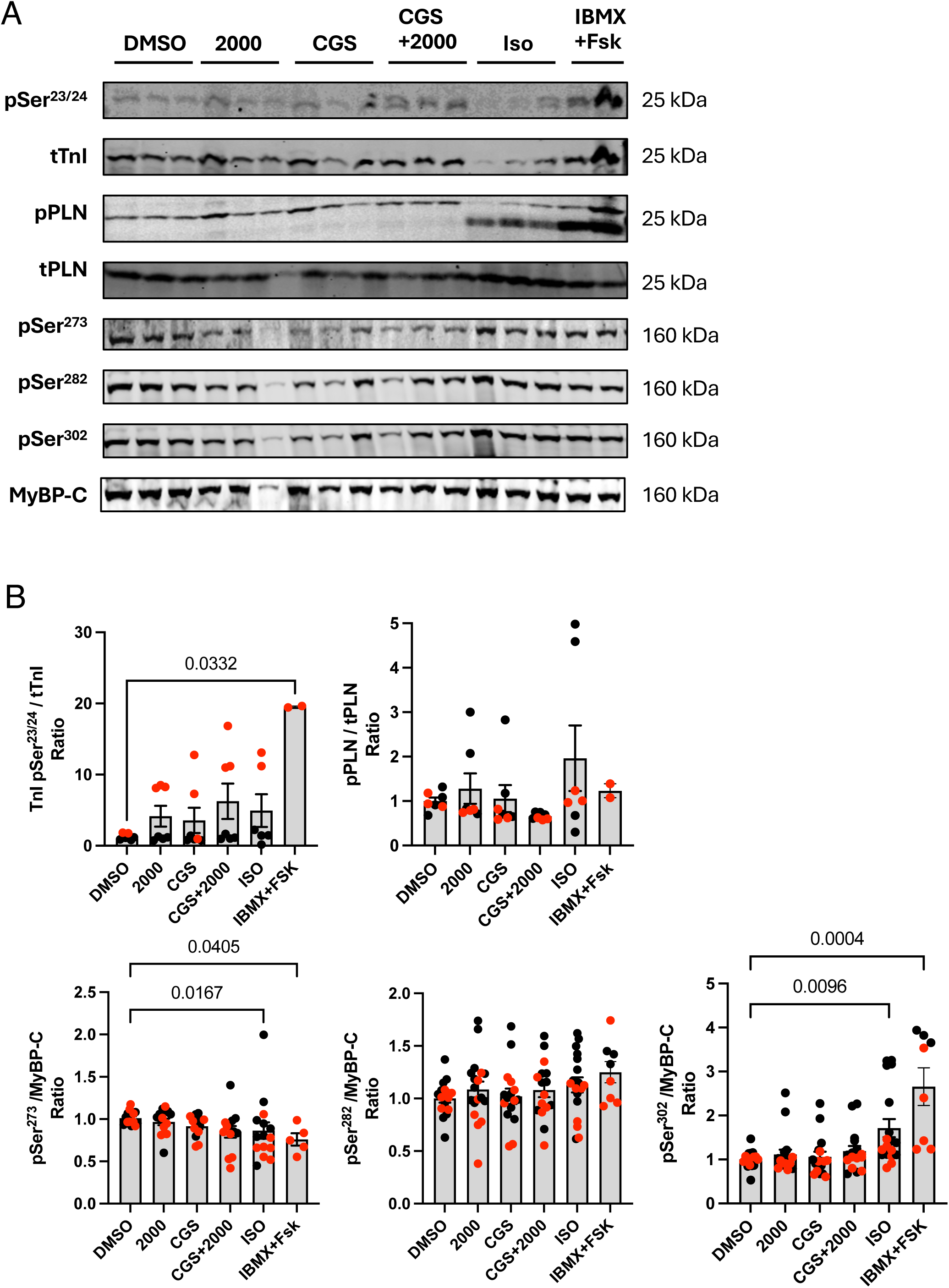
A) Representative western blots of PKA phosphorylation sites and total respective proteins, and B) quantitation of densitometry values compared to control (DMSO). Male and female data are shown as, respectively, black and red dots. Two-way ANOVA Kruskal-Wallis test with Dunn’s post-hoc analysis values shown (n=2-15, N=10).

**Supplemental Figure 6.**
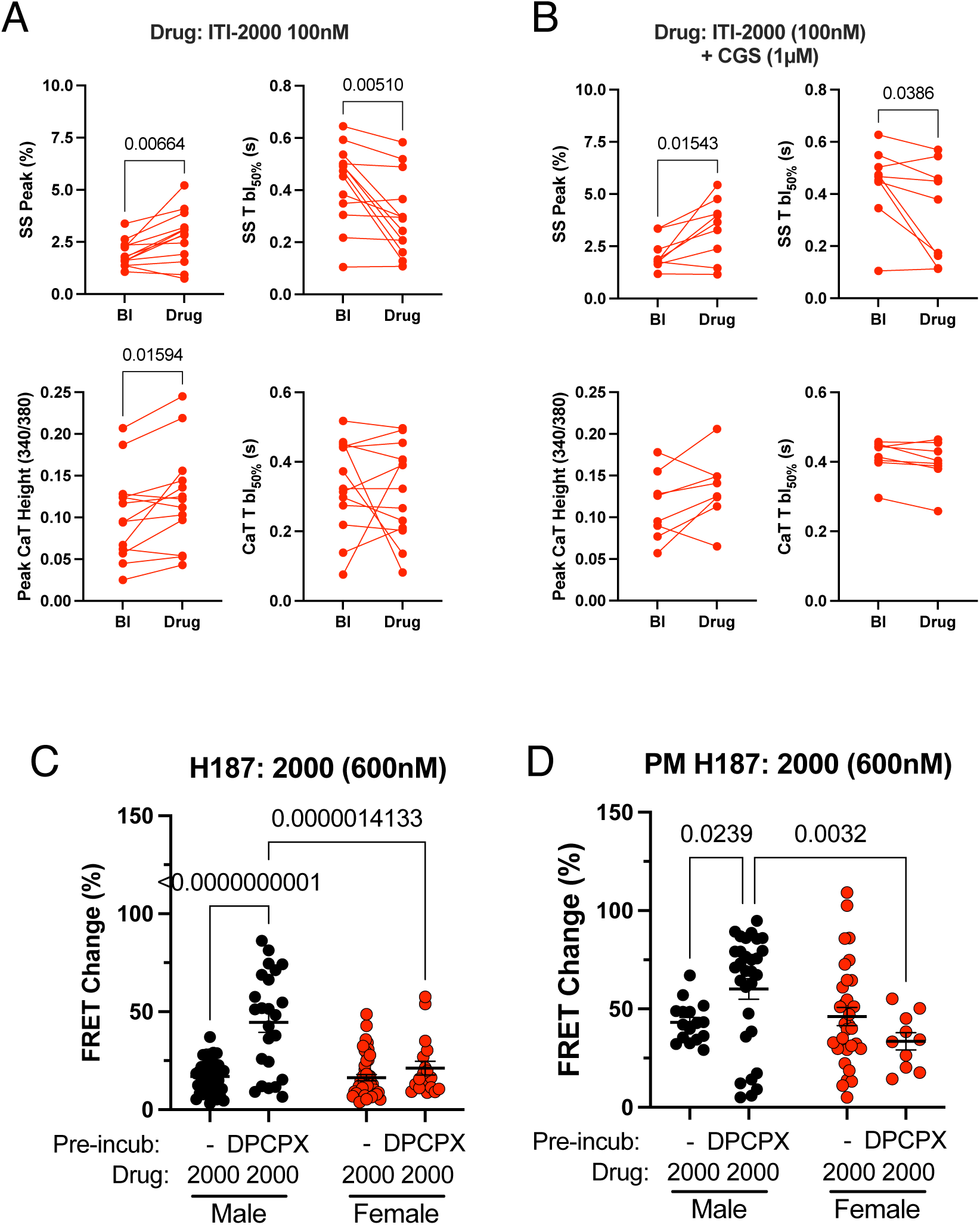
Cell SS and CaT changes upon A) 2000 (100nM) or B) 2000 (100nM) and CGS (1µM). Paired two-tailed Student’s t test values are shown (n=8-13, N=3-5). Cardiomyocytes expressing the C) cytosolic cAMP sensor H187 or D) plasma membrane cAMP sensor pm-H187 were treated with 2000 (600nM) with or without prior treatment with the A_1_R antagonist DPCPX (1µM). Group averaged FRET change values are plotted. Two-way ANOVA in C: F (1, 120) = 16.30, p=0.000096, n=18-44, N=4; and in D: F (1, 84) = 6.731, p=0.0112, n=10-32, N=4. Uncorrected Fisher’s LSD values post-hoc test results are shown.

**Supplemental Figure 7.**
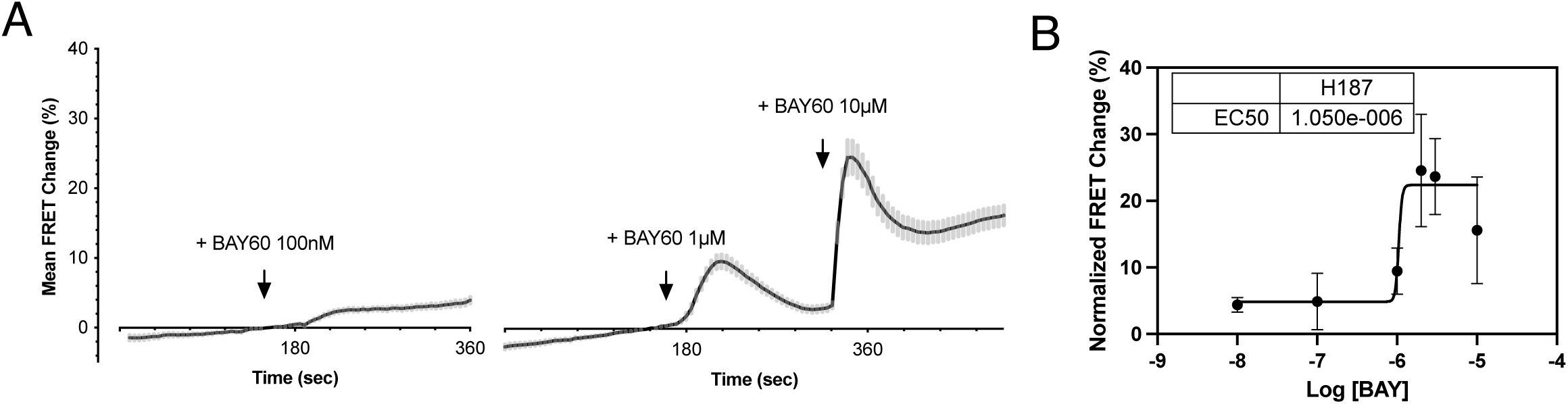
A) Guinea pig cardiomyocytes expressing the cytosolic cAMP sensor H187 were treated with various BAY60-6583 (BAY60) dose B) Log-transformed dose response curve of BAY60-6558 fitted with a four-parameter variable slope least squares fit (n=7-83, N=3).

**Supplemental Table 1.** IonOptix measurements in healthy male vs female guinea pigs.

|  |  |  | 2000 |  |  |  | CGS |  |  |  | CGS+2000 |  |  |  |
| --- | --- | --- | --- | --- | --- | --- | --- | --- | --- | --- | --- | --- | --- | --- |
|  |  |  | Baseline | n | Drug | n | Baseline | n | Drug | n | Baseline | n | Drug | n |
| Male | SS | Peak % | 2.60±0.24 | 22 | 2.35±0.35 | 22 | 2.30±0.25 | 24 | 2.93±0.42 | 24 | 2.67±0.26 | 27 | 4.63±0.62**** | 27 |
|  |  | Tpeak 50% (s) | 0.052±0.010 | 19 | 0.036±0.007**** | 19 | 0.055±0.007 | 23 | 0.040±0.007* | 23 | 0.025±0.003 | 22 | 0.020±0.002 | 22 |
|  |  | Tbl 50% (s) | 0.30±0.03 | 22 | 0.29±0.03 | 22 | 0.28±0.02 | 22 | 0.23±0.02*** | 22 | 0.19±0.02 | 23 | 0.14±0.02 | 23 |
|  | CaT | Peak (340/380) | 0.16±0.01 | 21 | 0.14±0.01 | 21 | 0.20±0.02 | 24 | 0.22±0.03 | 24 | 0.14±0.01 | 27 | 0.15±0.02 | 27 |
|  |  | Tpeak 50% (s) | 0.0113±0.0014 | 20 | 0.0090±0.0013** | 20 | 0.0141±0.0014 | 19 | 0.0104±0.0015*** | 19 | 0.0093±0.0012 | 24 | 0.0067±0.0008*** | 24 |
|  |  | Tbl 50% (s) | 0.31±0.03 | 21 | 0.31±0.03 | 21 | 0.26±0.02 | 21 | 0.23±0.02* | 21 | 0.21±0.02 | 25 | 0.21±0.02 | 25 |
| Female | SS | Peak % | 2.68±0.19 | 30 | 3.87±0.43*** | 30 | 2.55±0.15 | 36 | 2.76±0.31 | 36 | 2.67±0.15 | 48 | 3.47±0.32* | 48 |
|  |  | Tpeak 50% (s) | 0.050±0.004 | 29 | 0.041±0.003** | 29 | 0.051±0.006 | 31 | 0.042±0.004 | 31 | 0.049±0.003 | 42 | 0.035±0.002**** | 42 |
|  |  | Tbl 50% (s) | 0.40±0.03 | 30 | 0.30±0.03**** | 30 | 0.35±0.03 | 36 | 0.34±0.03 | 36 | 0.38±0.02 | 48 | 0.29±0.02**** | 48 |
|  | CaT | Peak (340/380) | 0.15±0.01 | 26 | 0.17±0.02* | 26 | 0.15±0.01 | 32 | 0.14±0.01 | 32 | 0.16±0.01 | 44 | 0.17±0.01 | 44 |
|  |  | Tpeak 50% (s) | 0.0127±0.0009 | 22 | 0.0129±0.0011 | 22 | 0.0091±0.0005 | 28 | 0.0096±0.0006 | 28 | 0.0102±0.0005 | 39 | 0.0095±0.0005 | 39 |
|  |  | Tbl 50% (s) | 0.44±0.02 | 25 | 0.39±0.01 | 25 | 0.37±0.02 | 32 | 0.37±0.02 | 32 | 0.38±0.01 | 44 | 0.34±0.01*** | 44 |

\*p<0.05, \*\*p<0.01, \*\*\*p<0.001, \*\*\*\*p<0.0001, two-tailed paired Student's t-test

**Supplemental Table 2.**
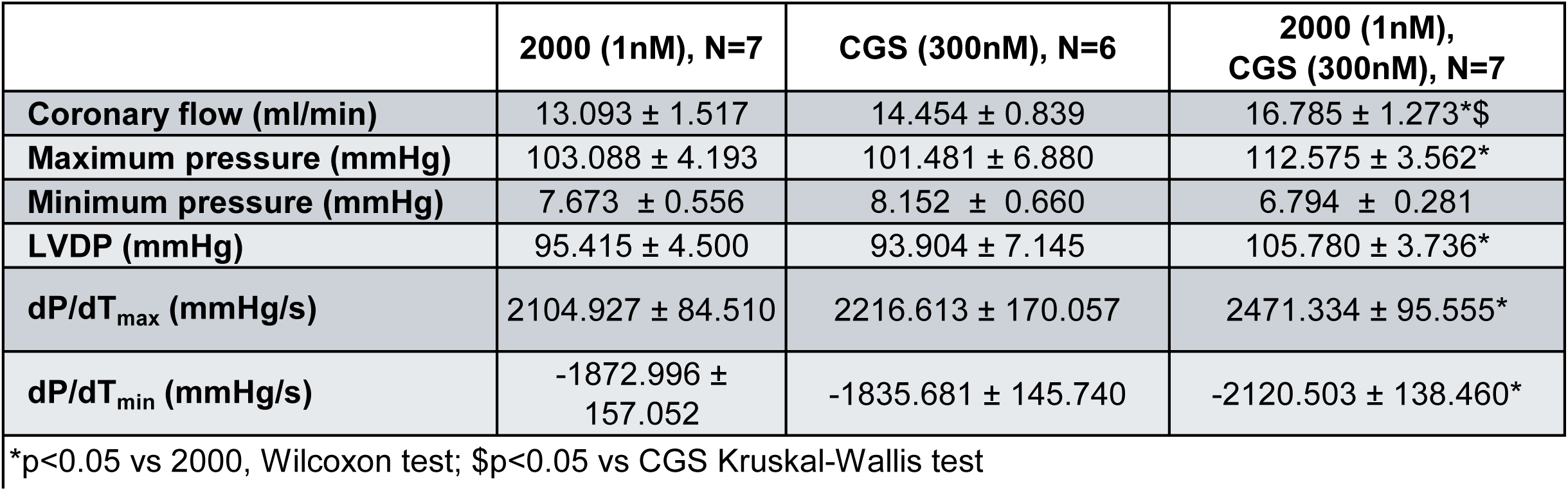
Ex vivo Langendorff measurements in guinea pigs.

**Supplemental Table 3.** BAY60-6583 or BAY+2000 response in healthy guinea pigs.

|  |  | BAY60-6583 |  |  |  | BAY+2000 |  |  |  |
| --- | --- | --- | --- | --- | --- | --- | --- | --- | --- |
|  |  | Baseline | n | Drug | n | Baseline | n | Drug | n |
| SS | Peak % | 2.4±0.1 | 74 | 1.8±0.13*** | 67 | 2.5±0.11 | 82 | 2.3±0.16 | 75 |
|  | Tpeak 50% (s) | 0.0484±0.0030 | 70 | 0.0360±0.0020*** | 65 | 0.0493±0.0030 | 71 | 0.0416±0.0024**** | 78 |
|  | Tbl 50% (s) | 0.35±0.014 | 74 | 0.31±0.014** | 74 | 0.38±0.014 | 81 | 0.32±0.015**** | 81 |
| CaT | Peak (340/380) | 0.14±0.00 | 73 | 0.13±0.00 | 73 | 0.15±0.00 | 78 | 0.14±0.00 | 76 |
|  | Tpeak 50% (s) | 0.011±0.0005 | 58 | 0.010±0.0004 | 58 | 0.012±0.0006 | 70 | 0.011±0.0005** | 72 |
|  | Tbl 50% (s) | 0.35±0.011 | 74 | 0.32±0.012* | 74 | 0.38±0.011 | 77 | 0.35±0.011*** | 77 |

\*\*p<0.01, \*\*\*p<0.001, \*\*\*\*p<0.0001, two-tailed paired Student's t-test

**Supplemental Table 4.** Echocardiogram Values in Guinea Pigs.

| **Supplemental Table 4. Echocardiogram Values in Guinea Pigs** |  |  |
| --- | --- | --- |
|  | **Sham (N=7)** | **AC (N=5)** |
| **BW (g)** | 661.4±57.34 | 698.6±78.03 |
| **HR (bpm)** | 255.6±10.08 | 257.0±13.85 |
| **Ejection Fraction (EF, %)** | 66.83±1.95 | 31.80±1.29* |
| **Fractional Shortening (FS, %)** | 38.37±1.50 | 15.88±0.62* |
| **LVID;d** | 8.23±0.24 | 9.88±0.9 |
| **LVAW;d** | 2.08±0.11 | 1.8±0.29 |
| **LVAW;s** | 3.28±0.18 | 2.25±0.24* |
| **LVPW;d** | 2.26±0.13 | 1.95±0.27 |
| **LVPW;s** | 3.19±0.15 | 2.21±0.25* |
| **Left Ventricle Weight (LVW)** | 1165±115.4 | 1451±98.2 |
| **LV/BW** | 1.843±0.03 | 2.143±0.16* |
| *p≤0.01 Mann-Whitney test |  |  |

**Supplemental Table 5.** IonOptix measurements in Sham guinea pigs.

|  |  | 2000 |  |  |  | CGS |  |  |  | CGS+2000 |  |  |  |
| --- | --- | --- | --- | --- | --- | --- | --- | --- | --- | --- | --- | --- | --- |
|  |  | Baseline | n | Drug | n | Baseline | n | Drug | n | Baseline | n | Drug | n |
| SS | Peak % | 2.24±0.2239 | 22 | 2.127±0.2496 | 22 | 1.930±0.158 | 25 | 2.660±0.3121* | 25 | 2.434±0.199 | 36 | 3.205±0.422* | 36 |
|  | Tpeak 50% (s) | 0.034±0.0035 | 21 | 0.027±0.0021* | 21 | 0.036±0.0039 | 24 | 0.031±0.0027 | 24 | 0.048±0.0049 | 36 | 0.040±0.0031* | 36 |
|  | Tbl 50% (s) | 0.358±0.03 | 22 | 0.269±0.02*** | 22 | 0.30±0.03 | 25 | 0.27±0.02 | 25 | 0.35±0.02 | 36 | 0.33±0.02 | 36 |
| CaT | Peak (340/380) | 0.195±0.017 | 22 | 0.197±0.020 | 22 | 0.176±0.015 | 24 | 0.195±0.019 | 24 | 0.231±0.018 | 34 | 0.238±0.019 | 34 |
|  | Tpeak 50% (s) | 0.009±0.0010 | 19 | 0.008±0.0005 | 21 | 0.010±0.0007 | 20 | 0.011±0.0008 | 20 | 0.013±0.0009 | 34 | 0.013±0.0009 | 34 |
|  | Tbl 50% (s) | 0.203±0.029 | 22 | 0.169±0.022* | 22 | 0.269±0.026 | 24 | 0.271±0.017 | 24 | 0.217±0.021 | 34 | 0.201±0.017 | 34 |
|  |  | BAY |  |  |  | BAY+2000 |  |  |  |  |  |  |  |
|  |  | Baseline | n | Drug | n | Baseline | n | Drug | n |  |  |  |  |
| SS | Peak % | 2.320±0.2229 | 30 | 2.300±0.2573 | 30 | 2.222±0.215 | 19 | 1.817±0.360 | 19 |  |  |  |  |
|  | Tpeak 50% (s) | 0.035±0.0026 | 30 | 0.033±0.0028 | 30 | 0.037±0.0039 | 18 | 0.035±0.0032 | 18 |  |  |  |  |
|  | Tbl 50% (s) | 0.31±0.03 | 30 | 0.29±0.02 | 30 | 0.39±0.05 | 19 | 0.33±0.03 | 19 |  |  |  |  |
| CaT | Peak (340/380) | 0.18±0.015 | 30 | 0.18±0.017 | 30 | 0.163±0.018 | 19 | 0.137±0.014* | 19 |  |  |  |  |
|  | Tpeak 50% (s) | 0.009±0.0007 | 30 | 0.010±0.0008 | 30 | 0.012±0.0016 | 17 | 0.010±0.0010 | 17 |  |  |  |  |
|  | Tbl 50% (s) | 0.230±0.017 | 30 | 0.203±0.018 | 30 | 0.217±0.027 | 18 | 0.219±0.025 | 18 |  |  |  |  |

\*p<0.05, \*\*p=0.0003 paired two-tailed Student's t-test

**Supplemental Table 6.**
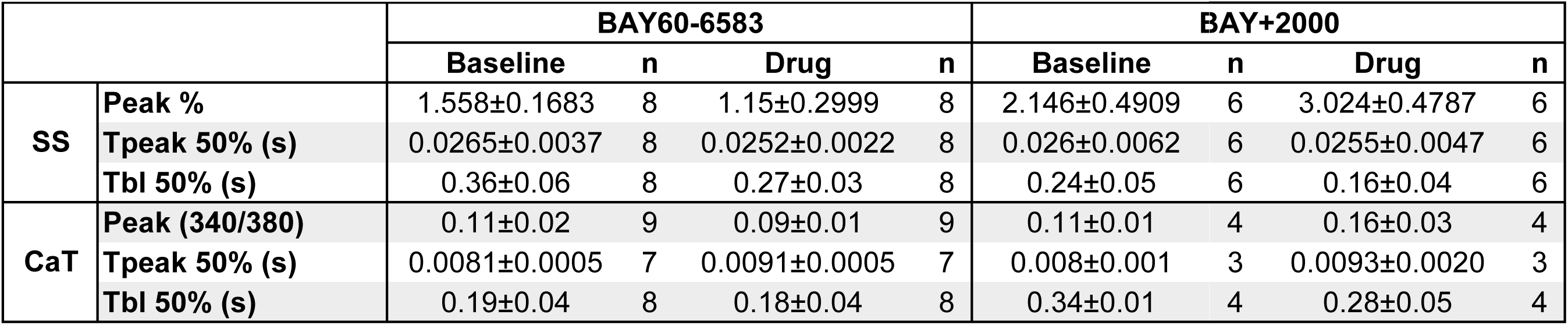
BAY60-6583 or BAY+2000 response in guinea pigs with HF.

## Notes

### Competing Interest Statement

The authors have declared no competing interest.

